# Global Analysis of the Mammalian MHC class I Immunopeptidome at the Organism-Wide Scale

**DOI:** 10.1101/2020.09.28.317750

**Authors:** Peter Kubiniok, Ana Marcu, Leon Bichmann, Leon Kuchenbecker, Heiko Schuster, David Hamelin, Jérome Despault, Kevin Kovalchik, Laura Wessling, Oliver Kohlbacher, Stefan Stevanovic, Hans-Georg Rammensee, Marian C. Neidert, Isabelle Sirois, Etienne Caron

**Affiliations:** CHU Sainte-Justine Research Center, Montreal, QC H3T 1C5, Canada; Department of Immunology, Interfaculty Institute for Cell Biology, University of Tübingen, Tübingen, Baden-Württemberg, 72076, Germany; Cluster of Excellence iFIT (EXC 2180) “Image-Guided and Functionally Instructed Tumor Therapies”, University of Tübingen, Tübingen, Baden-Württemberg, 72076, Germany; Applied Bioinformatics, Dept. of Computer Science, University of Tübingen, Tübingen, Baden-Württemberg, 72074, Germany; Immatics Biotechnologies GmbH, Tübingen, 72076, Baden-Württemberg, Germany; DKFZ Partner Site Tübingen, German Cancer Consortium (DKTK), Tübingen, Baden-Württemberg, 72076, Germany; Institute for Bioinformatics and Medical Informatics, University of Tübingen, Tübingen, Baden-Württemberg, 72076, Germany; Biomolecular Interactions, Max Planck Institute for Developmental Biology, Tübingen, Baden-Württemberg, 72076 Germany; Cluster of Excellence Machine Learning in the Sciences (EXC 2064), University of Tübingen, Tübingen, Baden-Württemberg, PLZ, Germany; Translational Bioinformatics, University Hospital Tübingen, Tübingen, Baden-Württemberg, 72076 Tübingen, Germany; Clinical Neuroscience Center and Department of Neurosurgery, University Hospital and University of Zürich, Zürich, Zürich, 8057&8091, Switzerland; Department of Pathology and Cellular Biology, Faculty of Medicine, Université de Montréal, QC H3T 1J4, Canada

**Author notes:** Corresponding and Leading author: Etienne Caron. Equal contribution to this work (Peter Kubiniok, Ana Marcu, Leon Bichmann).

## Abstract

Understanding the molecular principles that govern the composition of the mammalian MHC-I immunopeptidome (MHC-Ii) across different primary tissues is fundamentally important to predict how T cell respond in different contexts *in vivo*. Here, we performed a global analysis of the mammalian MHC-Ii from 29 and 19 primary human and mouse tissues, respectively. First, we observed that different HLA-A, -B and -C allotypes do not contribute evenly to the global composition of the MHC-Ii across multiple human tissues. Second, we found that peptides that are presented in a tissue-dependent and -independent manner share very distinct properties. Third, we discovered that proteins that are evolutionarily hyperconserved represent the primary source of the MHC-Ii at the organism-wide scale. Finally, we uncovered new components of the antigen processing and presentation network that may drive the high level of heterogeneity of the MHC-Ii across different tissues in mammals. This study opens up new avenues toward a system-wide understanding of antigen presentation *in vivo* and may serve as ground work to understand tissue-dependent T cell responses in autoimmunity, infectious diseases and cancer.

## INTRODUCTION

In adaptive immunity, CD8+ T cells have the ability to eradicate abnormal cells through recognition of small peptide fragments presented by MHC (human leucocyte antigen (HLA) in humans) class I molecules. In this context, jawed vertebrates evolved an important antigen processing and presentation system capable of presenting thousands of different MHC class I peptides on the surface of virtually any nucleated cells (Neefjes et al., 2011). In mammals, around 200 different cell types are decorated by large repertoires of self-MHC-I-associated peptides, collectively referred to as the mammalian MHC-I immunopeptidome (MHC-Ii) (Caron et al., 2017; Vizcaíno et al., 2019).

The inter- and intra-individual complexity of the MHC-Ii account for its overall heterogeneity (Gfeller and Bassani-Sternberg, 2018; Maccari et al., 2017; Vizcaíno et al., 2019). In fact, each MHC-I allotype generally presents a distinct subset of peptide antigens, which are characterized by the presence of specific anchor residues that are necessary to bind MHC-I (Falk et al., 1991). In human, up to six different HLA-I allotypes are expressed at the individual level, and thousands, if not millions of different HLA-I allotypes are expressed across human populations, hence increasing enormously the inter-individual heterogeneity of the MHC-Ii (Robinson et al., 2017). In contrast, the mouse MHC-Ii is relatively simpler. For instance, in the C57BL/6 mouse strain, peptide antigens are presented by only two classical MHC-I molecules (H2D^b^ and H2K^b^), and a total of approximately 200 different MHC-I allotypes are expressed among the most commonly used mouse strains (http://www.imgt.org/IMGTrepertoireMH/Polymorphism/haplotypes/mouse/MHC/Mu_haplotypes.html). In addition to its allotype-dependent composition, the mammalian MHC-Ii is also complicated by its tissue-dependency. In fact, two pioneering mapping studies recently pointed toward large variations in the repertoire of MHC-I-associated peptides across different tissues (Marcu et al., 2021; Schuster et al., 2018). However, very little is known about the molecular principles that shape the tissue-dependent processing and presentation of peptide antigens at the organism level.

Classical biochemistry approaches have established the blueprint of antigen processing and presentation (Neefjes et al., 2011; Yewdell et al., 2003). In a nutshell, the biogenesis of peptides presented by MHC-I molecules is initiated with the transcription and translation of the source genes, and the resulting proteins are typically degraded by the proteasome and/or immunoproteasome in the nucleus and cytosol (Kincaid et al., 2011). Cytosolic peptides are rapidly targeted by cytosolic aminopeptidases, such as thimet oligopeptidase (TOP) (York et al., 2003), leucine aminopeptidase (LAP) (Towne et al., 2005), and tripeptidyl peptidase II (TPPII) (Reits et al., 2004), which trim and destroy most peptides. A fraction of peptides escapes destruction by translocation into the endoplasmic reticulum (ER) lumen via transporter associated with antigen presentation (TAP) (Reits et al., 2000; Yewdell et al., 2003). In the ER, peptides may be further trimmed by ER aminopeptidase associated with antigen processing (ERAAP) and then bind MHC-I molecules for stabilization by the peptide loading complex (Blees et al., 2017; Serwold et al., 2002). Once stable, MHC-I-peptide complexes are released from the ER and are transported to the cell surface for peptide presentation to CD8+ T cells.

Modern immunopeptidomics is driven by high-resolution mass spectrometry (MS) and investigates the composition and dynamics of the MHC-Ii (Caron et al., 2015a, 2015b). Complementing classical biochemistry techniques, immunopeptidomic technology platforms have yielded important systematic insights into the biogenesis of the MHC-Ii (Granados et al., 2015). For instance, they have refined binding motifs for a wide range of MHC-I alleles in human (Abelin et al., 2017; Gfeller and Bassani-Sternberg, 2018), they have indicated that large numbers of MHC-I peptides derive from genomic ‘hotspots’ (Müller et al., 2017; Pearson et al., 2016; Marcu et al. 2021) as well as non-coding genomic regions (Laumont et al., 2018), and they have demonstrated that abundant transcripts and proteins contribute preferentially to the composition of the MHC-Ii (Abelin et al., 2017; Bassani-Sternberg et al., 2015; Fortier et al., 2008; Granados et al., 2012; Pearson et al., 2016). Furthermore, immunopeptidomic approaches have validated that defective ribosomal products (DRiPs), immunoproteasome subunits as wells as other key players involved in the processing of peptide antigens (e.g. proteasome, ERAAP) markedly influence the repertoire of peptides presented by MHC-I molecules (Bourdetsky et al., 2014; Milner et al., 2013; Nagarajan et al., 2016; Trentini et al., 2020; Verteuil et al., 2010).

The understanding of how the MHC-Ii is generated in different primary tissues *in vivo*, in human as well as in animal models, is fundamentally important to rationalize and predict how T cells respond in various contexts (Tscharke et al., 2015). However, immunopeptidomics studies that focused on the systematic deciphering of the MHC-Ii biogenesis have been almost exclusively conducted in transformed cells. Therefore, the rules that govern the composition and tissue-dependency of the mammalian MHC-Ii remains poorly understood and many fundamental questions remain unanswered to date. For instance, what is the relative contribution of individual HLA-I allotypes to the composition of the MHC-Ii within and across tissues? To what extent does the MHC-Ii conceal tissue-specific patterns/signatures that are conserved across species? What are the many transcription factors, proteases, amino- and carboxypeptidases involved in the generation and processing of MHC-I peptides in different tissues, and how does the expression and activity of those proteins influence the tissue-dependency and overall heterogeneity of the MHC-Ii at the organism-wide scale? In this study, we applied a systems-level, cross-species approach to tackle these fundamental questions.

## RESULTS

Two immunopeptidomic mapping studies have very recently drafted the first tissue-based atlases of the mouse and human MHC-Ii (Marcu et al., 2021; Schuster et al., 2018). These pioneering mapping efforts provide qualitative and semi-quantitative information about the currently detectable repertoire of MHC-I peptides in most organs, both in mouse and human. Specifically, the mouse atlas was generated from 19 normal primary tissues extracted from C57BL/6 mice expressing H2K^b^ and H2D^b^ (Schuster et al., 2018). The human atlas was generated from 29 human benign tissues extracted from 21 different subjects expressing a total of 51 different HLA-I allotypes (Marcu et al., 2021) (**Figure 1**). Those HLA-I allotypes cover the most frequent HLA-A, -B and -C alleles in the world. Below, we first focused on the analysis of the MHC-Ii in different mouse and human tissues to provide a general understanding of the heterogenicity, tissue-dependency and conservation patterns of the MHC-Ii. Next, we connected tissue immunopeptidomes to RNA-seq and protein expression data found in various tissue-based atlases (Geiger et al., 2013; Söllner et al., 2017; Wang et al., 2019) to dissect how the mammalian MHC-Ii is being shaped in different tissues (**Figure 1**).

**Figure 1.**
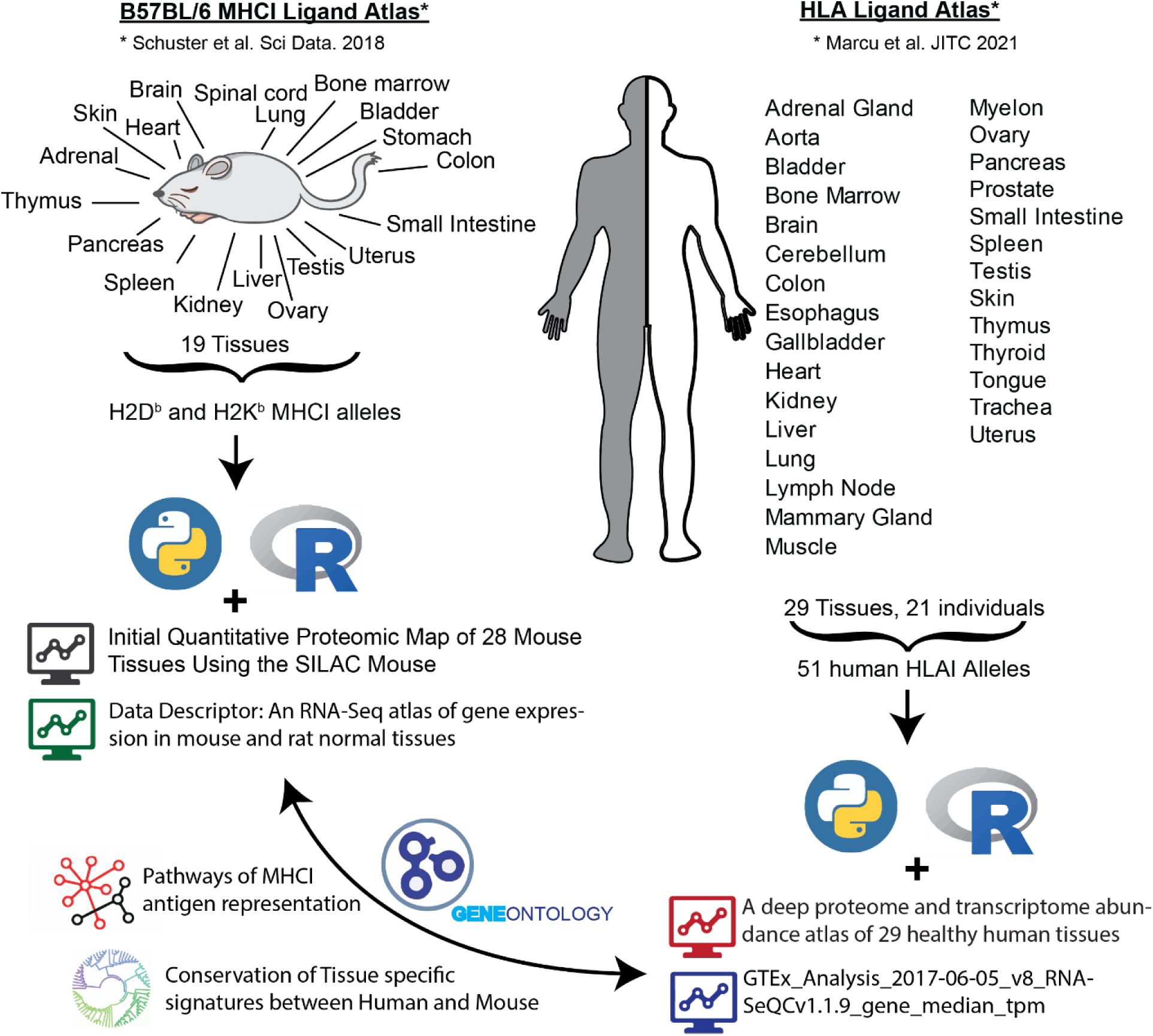
Overview of immunopeptidomics, proteomics and transcriptomics datasets analyzed. (**Left hand side**) Graphic description of the Mouse B57BL/6 MHCI Ligand Atlas, which was connected with two published proteomics and mRNA expression atlases of mouse tissues. (**Right hand side**) Graphic description of the Human HLAI Ligand Atlas, which was connected with two published proteomics and mRNA expression atlases of human tissues.

### HLA-I allotypes are unevenly represented across tissue immunopeptidomes

A key open question regarding the heterogeneity of the human MHC-Ii is whether individual HLA-I allotypes contribute evenly or unevenly to the composition of the MHC-Ii across different tissues. In fact, every subject presents up to two HLA-A, two HLA-B and two HLA-C allotypes. If all allotypes were evenly represented at the cell surface across tissues, one would expect similar proportions of peptides assigned to each allotype in all tissues. To address this question, we first assessed the global tissue distribution of all detectable peptides that were assigned to HLA-A, -B and -C. Among 29 sampled benign tissues extracted from a total of 21 different subjects, we found HLA-A, -B and -C immunopeptidomes to be unevenly represented across tissues (**Figure 2A**). To increase the resolution of this analysis, we investigated the contribution of each HLA-A, -B and - C allotypes expressed in the three subjects for which the most tissues had been sampled (i.e. AUT-DN11, AUT-DN13 and AUT-DN12) (**Figure 2 B-D**). Consistently, we found differential peptide distributions across tissues for many HLA-I allotypes. For instance, ∼55% of peptides in the Colon of subject AUT-DN12 were assigned to A*02:01 compared to ∼22% on average in all other tissues, resulting in an enrichment of about 2.5-fold for A*02:01 (**Figure 2D**). The enrichment of A*02:01 peptides in the Colon of subject AUT-DN12 was also further accompanied by an under-representation of A*11:01, B*15:01 and B*35:01 in the Colon, and an enrichment of C*03:04 and C*04:01 alleles (**Figure 2D**). Similarly, we also noted that ∼50% of peptides in the liver of subject AUT-DN13 were assigned to HLA-B40:02 compared to ∼20% on average in all other tissues, resulting in an enrichment of about 2.5-fold for this specific HLA-B allotype in this particular subject (**Figure 2C**). To provide a global picture about enrichment values that are associated to individual HLA-I allotypes, we calculated the average enrichment of all HLA-I allotypes across the investigated subjects and highlighted alleles that were enriched by more than 1.5-fold in at least one tissue (**Figure 2E**). This analysis highlighted 37 enrichment values distributed across 27 specific HLA-I allotypes and 18 different tissues (**Figure 2E**). Overall, those enrichment values ranged from 1.5 to 8.1-fold, and nine (out of 16) HLA-A, 9 (out of 21) HLA-B and 9 (out of 14) HLA-C allotype were assigned in at least one tissue with an enrichment value above 1.5-fold. Thus, our results show that HLA-I allotypes do not contribute evenly to the composition of the MHC-Ii across different tissues and subjects, and therefore, considerably contribute to the overall heterogeneity of the human MHC-Ii.

**Figure 2.**
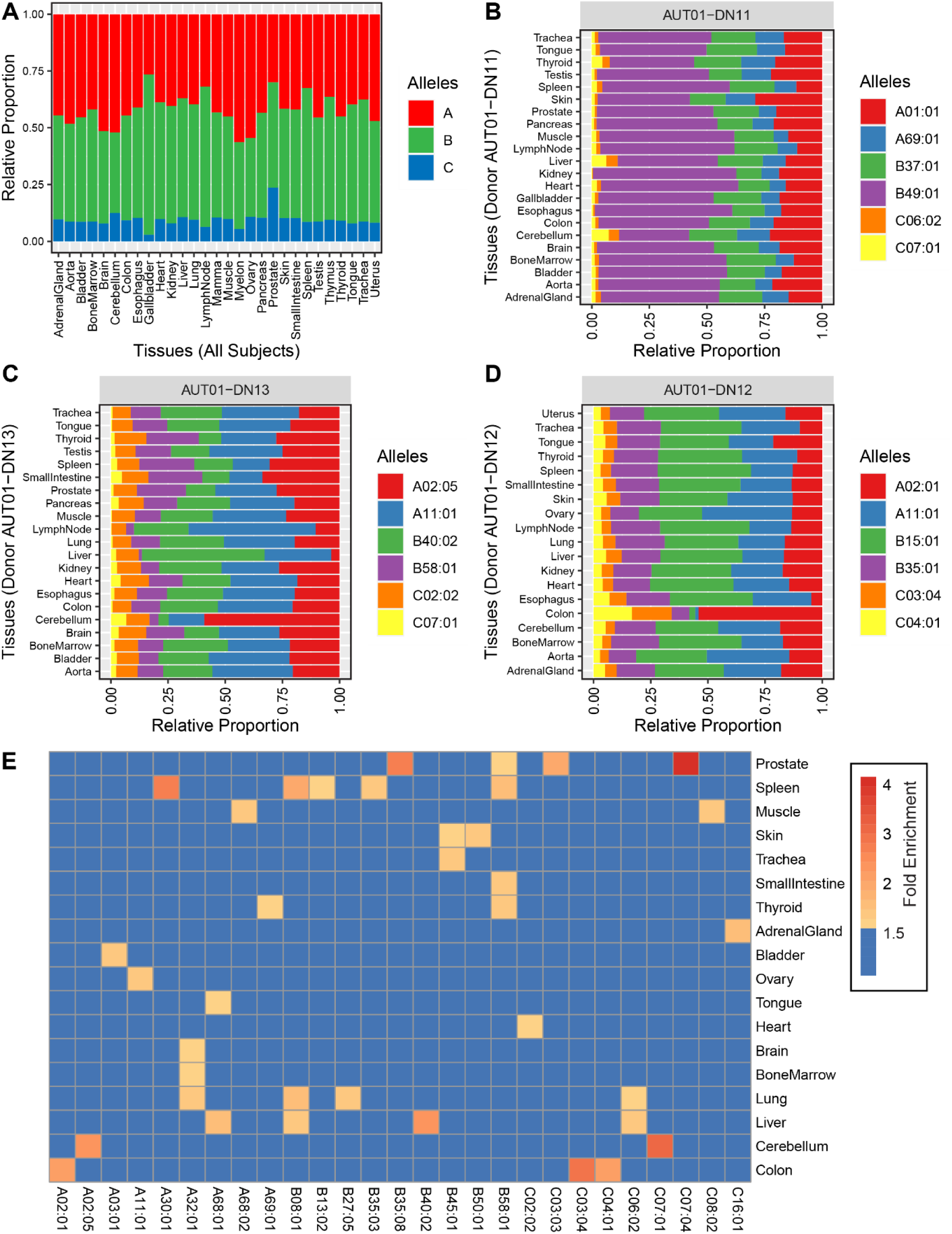
Distribution of HLA-A-, B- and C-specific immunopeptidomes across human tissues. **(A)** Relative proportion of individual HLA-A-, B- and C-specific immunopeptidomes per tissue among all subjects. **(B-D)** Relative proportion of individual HLA allele-specific immunopeptidomes per tissue for AUT-DN11 (B), AUT-DN13 (C) and AUT-DN12 (D). **(E)** Enrichment of HLAI allotypes across all tissues sampled. Average enrichment values are depicted where allotypes were sampled across several subjects.

### The high level of heterogeneity among immunopeptidomes of different tissues shows pronounced similarities between mouse and human

Antigen processing and presentation is a conserved and ubiquitous biological process in mammals. Here, we hypothesized that the MHC-Ii of different tissues might conceal tissue-dependent immunopeptidomic patterns/signatures that are conserved between mouse and human. First, we looked at the distribution of MHC-I peptide counts that were detected by MS across different mouse (**Figure 3A**) and human (**Figure 3B**) tissues. Expectedly, we noted that specific mouse organs yielded high numbers of MHC-I peptides (e.g. Spleen) whereas immune privilege organs (e.g., Brain and Testis) yielded low numbers of MHC-I peptides (**Figure 3A**). Very similar observations were made in human (**Figure 3B**) (Marcu et al., 2021). In fact, direct comparison of MHC-I peptide counts between mouse and human tissues resulted in a positive correlation (R-squared value = 0.44) (**Figure 3C**). Next, we performed principal component analyses (PCA) of tissue dependent intensities of mouse and human MHC-I peptides (**Figure 3D,E**). PCA were performed from highly heterogenous immunopeptidomic data integrating peptides and corresponding intensities presented by 2 mouse and 51 human MHC-I allotypes, respectively. Despite the high heterogeneity, our analysis revealed two main clusters in each species. Notably, immune-related organs clustered together in both species (see cluster 1 in **Figure 3D,E**). Immune clusters included Spleen, Bone Marrow, Lymph nodes and Thymus (mouse), as well as other types of non-immune related organs such as Kidney, Lung, Liver and Colon. The described observations raised the following question: what are the MHC-I peptides that are either shared or unique across these tissues?

**Figure 3.**
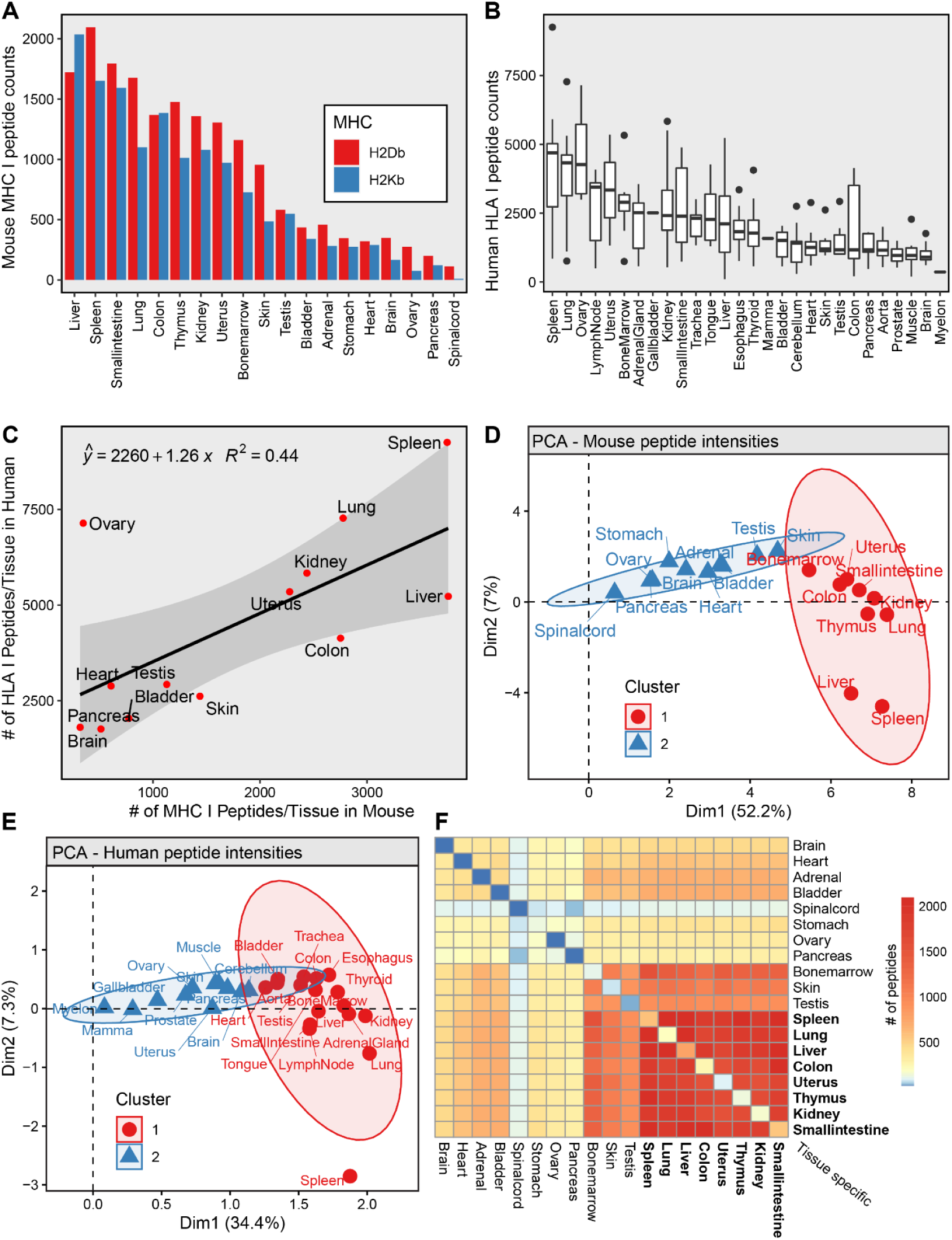
Comparison of tissue dependent MHCI- (Mouse) and HLAI (Human) -associated peptides. **(A)** MHCI peptide counts for each sampled mouse tissue, colors depict the MHCI alleles (H2D^b^ and H2K^b^). **(B)** HLAI peptide counts for all sampled human tissues. Boxplots are represented as several tissues were sampled across different individuals. **(C)** Comparison of MHCI peptide counts/tissue (Mouse) and HLAI peptide counts/tissue (Human). **(D)** Principal component analysis of the measured intensities (log10) of MHCI peptides (Mouse). **(E)** Principal component analysis of the measured intensities (log10) of HLAI peptides (Human). **(F)** Tissue connectivity map of the ‘B57BL/6 MHCI Ligand atlas. Heatmap depicts the number of shared MHCI peptides across tissues (Mouse). Note: The number of uniquely observed/tissue-specific peptides can be found along the diagonal.

To address the above question, we created connectivity matrices, which summarize the number of MHC-I peptides shared and uniquely observed between all possible pairs of tissues in mouse (**Figure 3F**) and human (**Supplementary Figure 1**). The number of uniquely observed/tissue-specific peptides can be found along the diagonal of the connectivity matrices in **Figure 3F** and **Supplementary Figure 1**. In mouse, we observed that 13% (961 out of the 7665 unique peptides found in mouse) of the total H2D^b^/K^b^ immunopeptidome was shared across Spleen, Bone Marrow, Kidney, Lung, Liver and Colon (**Figure 3F**). As an example, 1881 peptides (25% of the total H2D^b^/K^b^ immunopeptidome) were shared between Spleen and Kidney, and 1381 peptides (18% of the total H2D^b^/K^b^ immunopeptidome) were shared between Bone marrow and Liver (**Figure 3A**). In human, we observed that 4% of the total HLA-ABC immunopeptidome was shared across these six organs for all subjects. Once deconvolved by allotype or subject, we observed that, on average, 3% (range: 0.5% HLA-C*07:04 — 9% HLA-A*01:01 and B49:01) and 0.8% (range: 0.4% AUT-DN08 — 1.3%, AUT-DN12) of HLA-I peptides were shared across these organs, respectively (**Supplementary Figure 2**). In contrast, larger fractions of MHC-I peptides were found to be uniquely observed in each species. Overall, 42% (3212 out of 7665 unique peptides) and 44% (32187 out of 73639 unique peptides) of the total H2D^b^/K^b^- and HLA-ABC-immunopeptidome were uniquely observed in specific tissues, respectively. These peptides are further referred to as tissue-specific peptides. Thus, our data show, using the currently available technology, that a significant proportion of MHC-I peptides are tissue-specific whereas a relatively smaller proportion of peptides are shared across various immune and non-immune organs, both in mouse and human. To our knowledge, this is the first time that estimates of tissue-specific and - nonspecific MHC-I peptides are reported at the organism level. These two categories of MHC-I peptides may show distinct properties or trends, and were further investigated below.

### MHC-I peptides shared across multiple tissues are highly abundant and strong MHC-I binders

To investigate the properties of tissue-specific peptides versus those that are presented across a wide range of tissues, we sought to assess the influence of peptide abundance and MHC binding affinity on tissue distribution. Hence, we plotted the frequency of peptide measurements across mouse and human tissues against their average abundance or predicted MHC-I/HLA-I binding affinity (NetMHCpan4.0 rank score) (**Supplementary Figure 3** for mouse and **Supplementary Figures 4 and 5** for human). The human dataset has to be viewed in a subject-specific manner as each subject presents its own repertoire of HLA-I alleles. In mouse, we found that increasing cross-tissue presentation of an MHC-I peptides strongly correlated with increasing peptide abundance and increasing affinity for the MHC-I molecules (decreasing NetMHCpan 4.0 rank score) (**Supplementary Figure 3**). The same behavior was generally observed in human, where peptides widely represented across tissues were highly abundant (**Supplementary Figure 4**) and predicted to be strong HLA-I binders in all subjects (**Supplementary Figure 5**). Together, these results show that peptide abundance and binding affinity for MHC-I molecules are key properties that contribute to the widespread or tissue-specific presentation of peptides in the mammalian MHC-Ii.

### Tissue-specific MHC-I peptides arise from genes that are almost uniquely expressed in the peptide-producing tissue

Expression of tissue-specific source proteins contributes to shaping the tissue-specificity of the human MHC-Ii (Marcu et al., 2021). Pioneering work in mice also proposed that transcriptomic signatures of thymic cells can be conveyed to the cell surface in the MHC-Ii (Fortier et al., 2008). However, how gene expression shapes the composition of the mouse MHC-Ii across many different tissues *in vivo* has never been reported to date. To address this, we first assigned every mouse MHC-I peptide found in the tissue draft atlas of the MHC-Ii to its source gene. Using an RNA-Seq atlas of gene expression in mouse normal tissues (Söllner et al., 2017), we next assessed the transcript abundance of the MHC-I peptide source genes in nine tissues for which mRNA expression data were available (i.e. Brain, Colon, Heart, Kidney, Liver, Pancreas, Small intestine, Stomach and Thymus) (**Figure 4**). By doing so, we found that genes coding for any detectable MHC-I peptides as well as for tissue-specific MHC-I peptides were more actively transcribed compared to genes that were not coding for any detectable MHC-I peptides (**Figure 4 A,B**).

**Figure 4.**
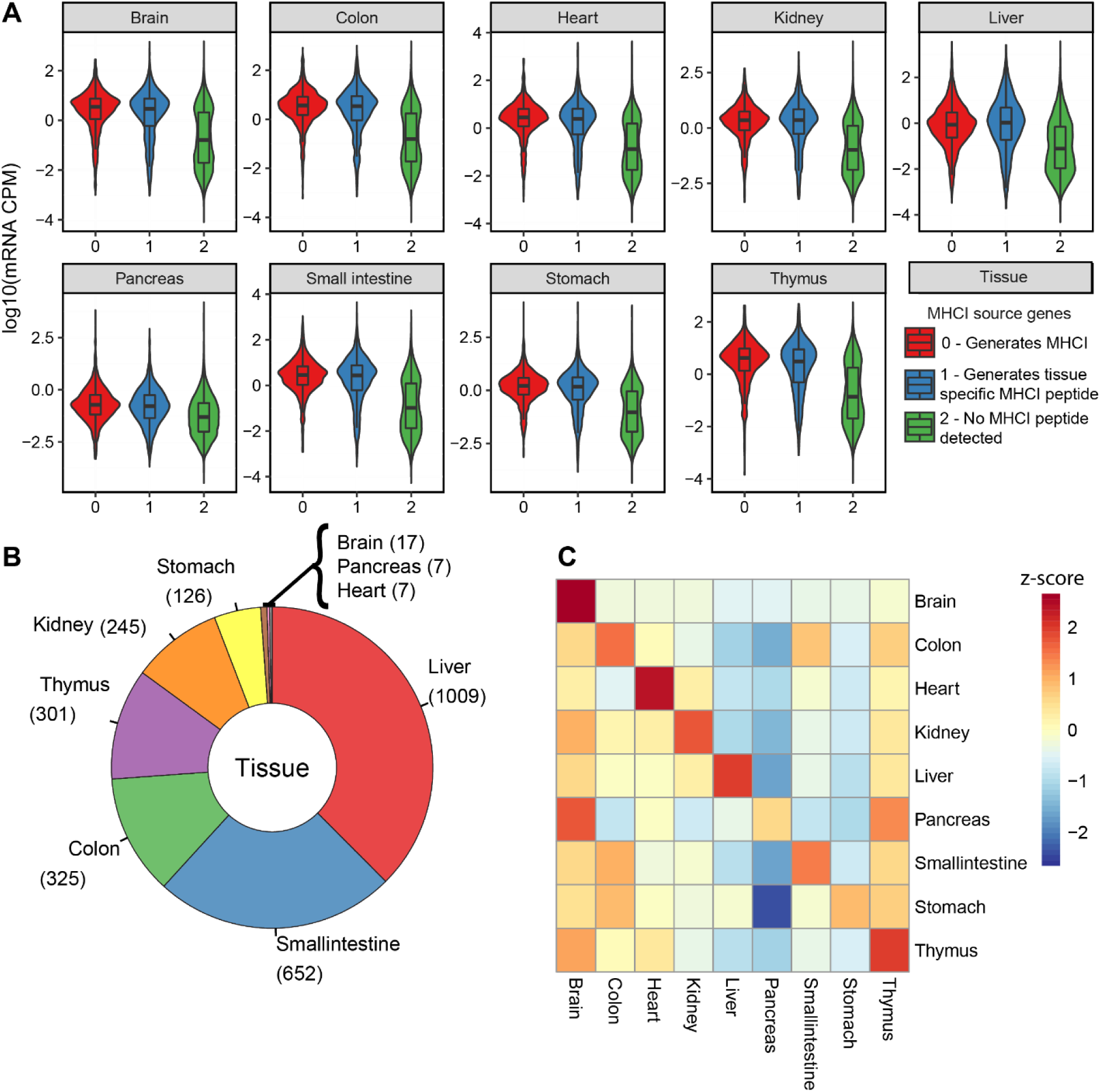
mRNA expression of MHCI source genes in multiple mouse organs. **(A)** Violin plots depicting the distribution of mRNA expression of genes which generate MHCI peptides (0), genes which generate tissue specific MHC peptides (1) or does not generate MHCI peptides (2). **(B)** Donut plot depicting the number of tissue-specific MHCI peptides found in tissues for which mRNA expression data is available (9 of 19 tissues sampled in the ‘B57BL/6 MHCI Ligand atlas’. **(C)** Heatmap representing the average mRNA expression of genes coding for tissue-specific MHCI peptides across tissues. Z-score is color coded.

Next, we reasoned that tissue-specific MHC-I peptides could derive from tissue-specific transcripts. To test this hypothesis, we averaged for every tissue the transcript abundance of genes coding for tissue-specific peptides and compared their expression across the nine tissues (**Figure 4C**). As depicted, we observed that brain-specific MHC-I peptides derived from genes that were uniquely expressed in the brain. Interestingly, liver-specific MHC-I peptides derived from genes that were predominantly, but not exclusively expressed in the liver—an expression pattern that was observed for seven out of nine tissues (colon, kidney, liver, heart, small intestine, stomach and thymus; **Figure 4C**). Thus, we provide new evidence at the organism-scale that tissue-specific MHC-I peptides are generally encoded from genes that are highly expressed in the same tissue of origin. Together, these results are in accordance with conclusions drawn in human (Marcu et al., 2021) and enforce the notion that gene expression plays a fundamental role in shaping the tissue specificity of the MHC-Ii in mammals.

### MHC-I peptides that are broadly presented across many tissues are encoded by genes that are highly expressed and evolutionarily hyperconserved

Above, we provided evidence that the MHC-Ii is composed of tissue-specific peptides as well as peptides that are widely presented across many different tissues. While tissue-specific MHC-I peptides appear to stem from genes predominantly expressed in the original tissue, we asked whether MHC-I peptides that were presented across most tissues derived from highly transcribed genes across the entire human or mouse genome. To answer this question, we created a selection of MHC-I peptides that were widely represented among the sampled tissues, referred herein as ‘housekeeping/universal MHC-I peptides’ (**Supplementary Figure 6A**). While this selection is straightforward for the mouse data where we considered peptides identified in all 19 of the 19 tissues as housekeeping/universal peptides, a more complex approach was needed to select those peptides in the human dataset where several subjects, each representing a specific set of HLA-I alleles, were present. Details about the selection of those peptides in the human immunopeptidome tissue draft are described in the methods section ‘Selection of Housekeeping/Universal Peptides’ and are visualized in **Supplementary Figure 6B-F** and **Supplementary Figure 7**). First, we found that the selected MHC-I peptides originated from 38 and 251 source genes in mouse and human, respectively (**Supplementary Tables 1 and 2,** and **Supplementary Figures 6 and 7**). Importantly, we discovered that these genes were among the most transcriptionally expressed genes across the entire mouse (**Figure 5A**) and human (**Figure 5B**) genome. This result is in line with the above observation that widely presented peptides across the organism are of high abundance (**Supplementary Figures 3 and 4**). Moreover, it is noteworthy that those housekeeping/universal MHC-I peptides did not preferentially originate from large (heavy) proteins, as it could have been expected due to the higher numbers of possible peptide antigen products from large proteins (**Supplementary Figure 8**).

**Figure 5.**
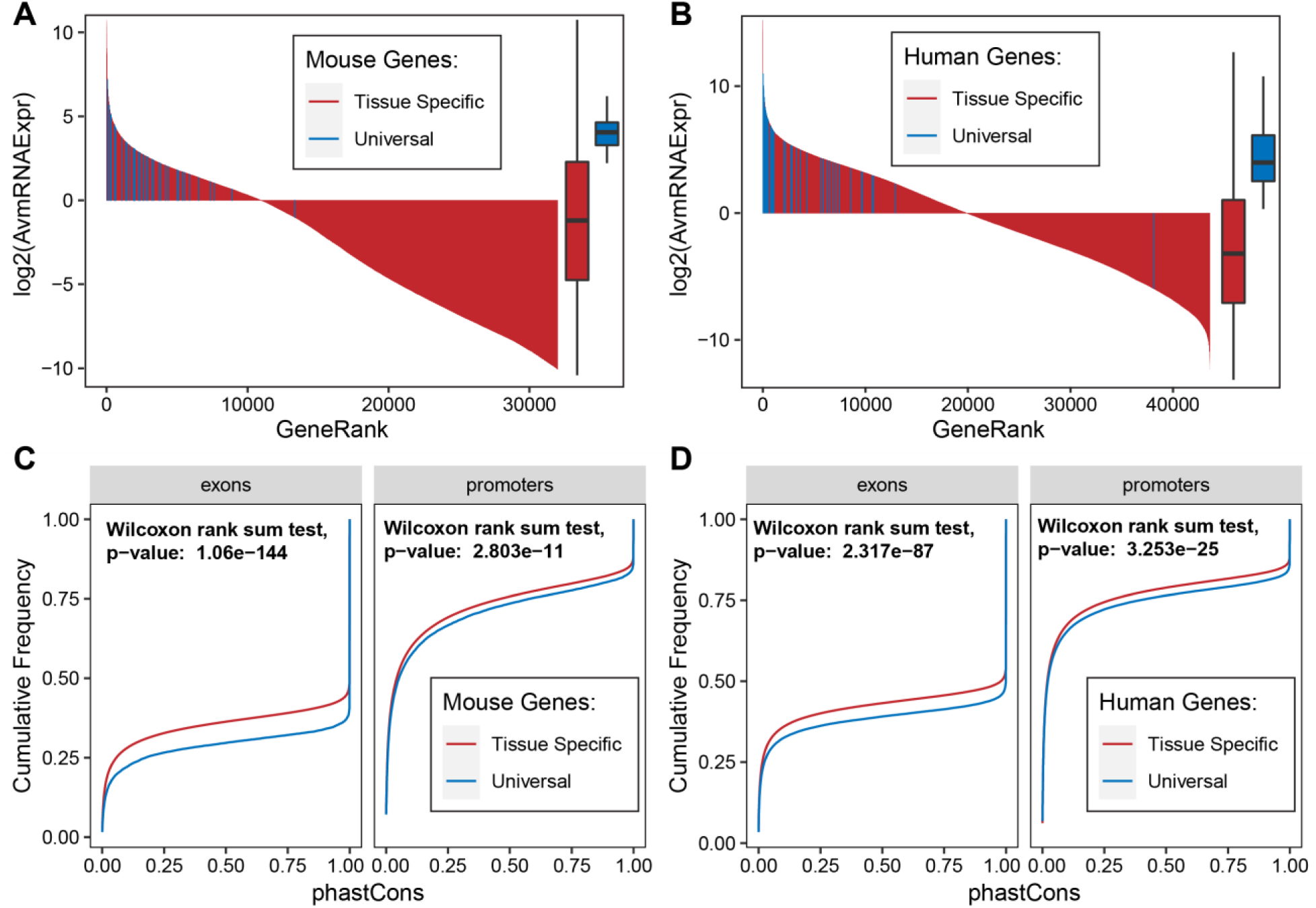
Expression and genetic conservation of genes coding for MHCI/HLAI peptides presented across most tissues (housekeeping/universal peptides). **(A,B)** mRNA expression of source genes of housekeeping/universal MHCI/HLAI peptides compared to all other mRNA transcripts in mouse (A) and human (B). **(C,D)** Exon and promoter conservation distributions of source genes of housekeeping/universal MHCI/HLAI peptides compared to source genes of tissue-specific MHCI peptides in mouse (C) and human (D).

Genes expressed in the majority of tissues in an organism play vital functions, are evolutionarily hyperconserved and are widely referred to as housekeeping genes (She et al., 2009; Zeng et al., 2016; Zhu et al., 2008). Akin to housekeeping genes, peptides that are represented in most tissues across an entire organism—referred above to as housekeeping/universal MHC-I peptides—could also originate from hyperconserved proteins as they may have co-evolved for millions of years with ancients and ubiquitous degradation systems to become the fundamental ground source of MHC-I peptides for most tissues. Hence, we hypothesized that universal MHC-I peptides are encoded by genes that are evolutionarily hyperconserved across evolution. To address this concept, we took advantage of the genome alignments between mouse and 59 vertebrates as well as between human and 99 vertebrates, made available by the UCSC Genome Browser (Lee et al., 2020) (see Materials and Methods section ‘Conservation of source genes from universal MHC-I peptides’). To assess evolutionarily conservation across species, PhastCons scores (Siepel et al., 2005), which predict the probability of conservation for every base-pair in the aligned genomes, were consulted for mouse and human genes of interest (see Materials and Methods for details). When comparing the conservation scores of tissue-specific MHC-I peptide source genes with those from housekeeping/universal MHC-I peptide source genes, the latter were significantly more conserved at the Promoter and Exon level, both in Mouse (p-value = 2.8×10^−11^; p-value = 1.06×10^−144^) (**Figure 5C**) and Human (p-value = 3.25×10^−25^; p-value = 2.32×10^−87^) (**Figure 5D**). For example, the conservation probability (PhastCons score) of 70% of the more conserved Exons (Cumulative Frequency > 0.3) of tissue-specific peptide source genes in mouse is greater than 20%, whereas the conservation probability of 70% of the more conserved Exons of housekeeping/universal peptide source genes in mouse is greater than 80%, making the latter set of genes preferentially hyperconserved. Thus, this analysis indicates that tissue-specific versus housekeeping/universal MHC-I peptide source genes do not share the same degree of conservation across evolution. Together, our results suggest, for the first time, that highly expressed and hyperconserved genes are preferential sources of MHC-I peptides at the organism-wide scale.

### Discovery of new components of the constitutive antigen processing and presentation network in mouse and human tissues

Differential expression and activity of antigen processing and presentation proteins across tissues may contribute to the observed variability in the composition of the MHC-Ii from one tissue to another (Rock et al., 2016). In this regard, transcript levels of HLA-I, TAP1/2 and immunoproteasome were very recently shown to correlate positively with the total number of MHC-I peptides detected across different human tissues (Marcu et al., 2021). To date, such correlative analysis has only been applied at the transcript level for a handful number of preselected immune-related genes and has never been performed at the protein level in a systematic fashion. Hence, we reasoned that an unbiased computational approach could be used to systematically identify any protein of the proteome for which their respective abundance across tissues correlates with the total number of MHC-I peptides across those same tissues. Therefore, we set out to apply this correlative approach at the proteome-wide scale using protein abundances measured across different mouse and human tissues from two tissue-based proteomics atlases generated by MS (**Figure 6A**) (Geiger et al., 2013; Wang et al., 2019).

**Figure 6.**
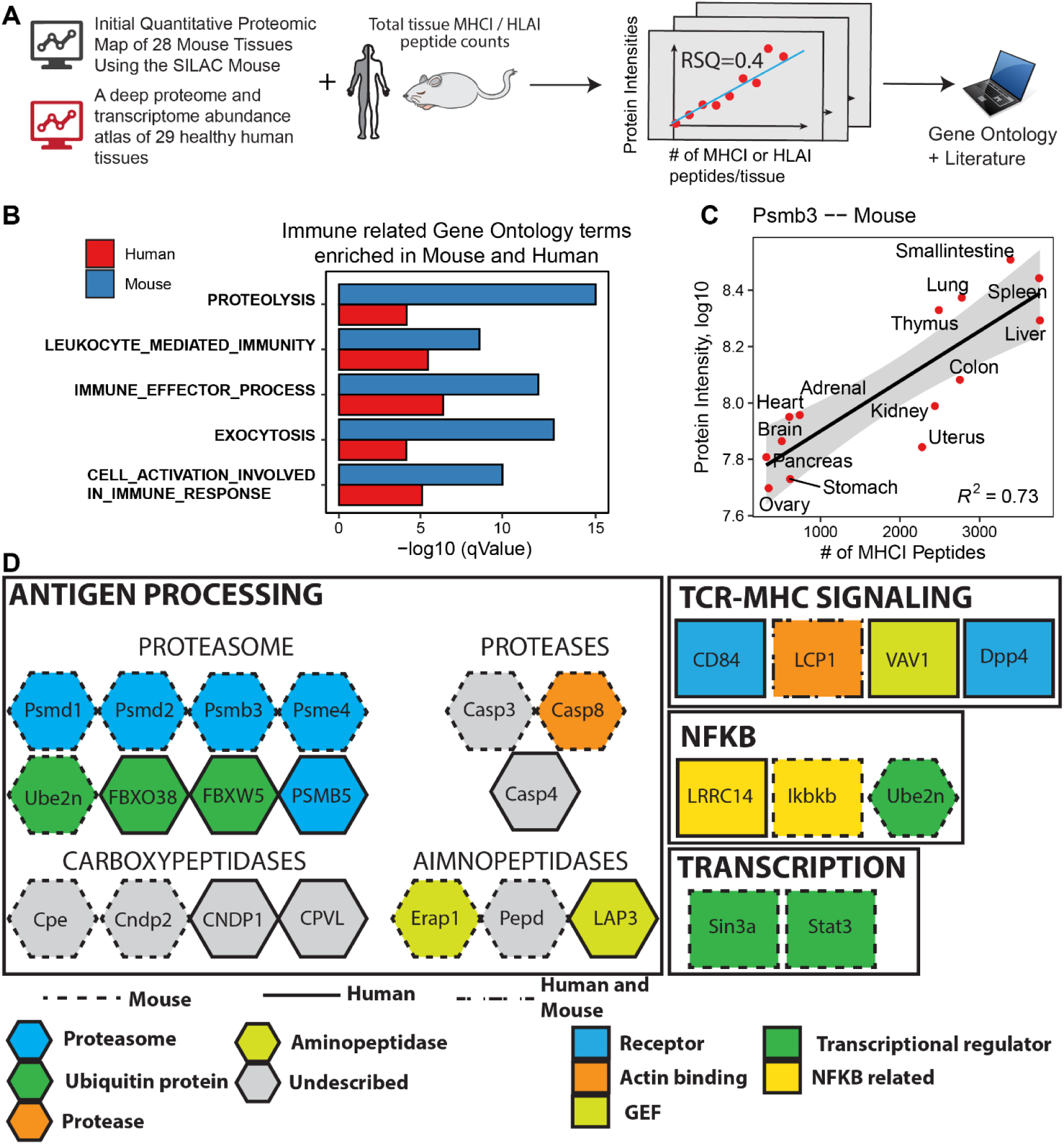
Correlation of protein abundances at the proteome-wide scale with the total number of MHCI or HLAI peptides detected across tissues. (**A**) Protein expression data from protein expression maps of mouse and human tissues were correlated with the total number of MHCI or HLAI peptides detected per tissue. Correlations were simulated for every protein measured across nine or more tissues. Significantly correlating proteins were further investigated. (**B**) Gene Ontology terms enriched from 164 mouse and 120 human proteins whose abundance significantly correlates with the number of MHCI/HLAI peptides counted per tissue. (**C**) Example correlation of proteasome subunit Psmb3 in mouse with MHCI peptides counted across tissues. (**D**) Protein modules identified from the global correlative analysis are associated to antigen generation, processing and recognition. Mouse and human proteins annotated to enriched GO terms were manually curated from the literature and were classified based on their respective biological function: proteasome, aminopeptidase, carboxypeptidase, protease, ubiquitin protein, guanine nucleotide–exchange factor (GEF), actin binding protein and transcriptional regulator and NFKB related. Proteins depicted in grey are uncharacterized enzymes of the antigen processing network.

First, we computed a total of 4,175 (on 4,175 gene coding proteins) and 70,656 (on 11,776 gene coding proteins) correlations in mouse and human, respectively (see Materials and Methods). Importantly, we found a subset of 164 and 120 correlating proteins in mouse and human, respectively, whose abundance significantly correlated with the total number of MHCI peptide counts in a given tissue (p-value < 0.01 and R-squared > 0.4 in Mouse; p-value < 0.05 and R-squared > 0.4 for at least two subjects in Human) (**Supplementary Figure 9A and B**, **Supplementary Tables 3 and 4**). From the 164 mouse proteins, 122 correlated positively (74%) and 42 correlated negatively (26%) with MHCI peptide counts. Out of the 120 significantly correlating human proteins, 74 correlated positively (62%) and 46 negatively (38%) **(Supplementary Figure 10).**

To broadly assess biological processes in which these proteins are implicated, we performed gene ontology (GO) analysis on these significantly correlating proteins (**Supplementary Table 5**). From the top 50 most significantly enriched GO terms implicated in biological processes in mouse and human, 15 were shared across both species (**Supplementary Figure 11**). Remarkably, the shared GO terms were attributed to proteins implicated in the regulation of the proteolysis, antigen processing and immune response (**Figure 6B,D**). A prominent example protein, PSMB3 whose tissue dependent intensity very well correlates with the amount of MHCI peptides found in each tissue is shown in **Figure 6C**. Furthermore, manual curation of the literature allowed us to associate those proteins to specific functional modules known to orchestrate transcription (e.g., STAT3, NFKB), TCR-MHC signaling (e.g., LCP1, VAV1) and antigen processing (e.g., PSMB3/5, PSME4, LAP3, ERAP1) (**Figure 6D**). Among the latter, many proteasome subunits, proteases, carboxy- and aminopeptidases were identified (**Figure 6D**). For example, PSMB3 is a component of the 20S core proteasome complex (Elenich et al., 1999; Huber et al., 2012); PSME4 is a proteasome activator subunit, also known as PA200 (Rêgo and Fonseca, 2019), and ERAP1 plays a central role in peptide trimming for the generation and presentation of MHC-I peptides (Serwold et al., 2002). For these three specific proteins, their abundance increased as a function of the number of MHC-I peptides (**Figure 6C**, **Supplementary Figure 12** and **Supplementary Tables 3, 4** and **5**). In contrast, we found the opposite trend for other proteins. For instance, abundance of Uchl1, Ube2n and PSMB5 proteins decreased as a function of the number of MHC-I peptides (**Supplementary Figure 10B, C and D**), the latter being known to be replaced by the immunoproteasome and thymoproteasome subunit PSMB8 and PSMB11 in immune and thymic cells, respectively (Murata et al., 2018). Most strikingly, we found four poorly characterized carboxypeptidases (CPE, CNDP1, CNDP2 and CPVL) showing significant correlations between protein abundance and number of MHC-I peptides across tissues (**Figure 6D**). This unexpected finding is interesting because very little is known about the role of carboxypeptidases in antigen processing. Further investigation is therefore required to determine the precise role of CPE, CNDP1, CNDP2 and CPVL in shaping the global composition of the mammalian MHC-Ii in health and diseases. Thus, our systems-level analysis allowed us to identify many known key players of the antigen processing network, thereby validating our computational approach, in addition to expand the network through identification of new components. Collectively, our study provides an unprecedented source of information regarding the biogenesis of the mammalian MHC-Ii and opens up new avenues to further explore the role of new proteolytic enzymes in antigen processing *in vivo*.

## DISCUSSION

The components of the antigen processing and presentation pathway shape how T cells respond to self and non-self (Rock et al., 2016). Those components have been traditionally discovered using hypothesis-driven approaches or genomic screening of cell lines presenting a phenotype of interest (Burr et al., 2019; Neefjes et al., 2011; Paul et al., 2011). MS-based immunopeptidomic approaches have also been used to validate the impact of those proteins on the global composition of the MHC-Ii using *in vitro* or *ex vivo* model systems (Alvarez-Navarro et al., 2015; Nagarajan et al., 2016; Verteuil et al., 2010). To date, no study has taken advantage of the uncharted combination of immunopeptidomic, proteomic, transcriptomic and genomic data from a range of different primary tissues to infer the fundamental principles that form the mammalian MHC-Ii. In fact, akin to systems immunology methods (Villani et al., 2018), we deployed in this study an unbiased immunopeptidomic data-driven strategy using multiple tissue-based omics datasets, both in mouse and human, to i) reinforce the notion that the composition of the mammalian MHC-Ii is highly context-dependent, ii) provide fundamental information about the tissue-dependency, conservation and biogenesis of the MHC-Ii at the organism-wide scale, and iii) uncover new proteins that may collectively orchestrate the content and tissue-specificity of the MHC-Ii.

In this study, we found that many proteins of the ubiquitin-proteasome degradation system as well as many proteases, amino- and carboxypeptidases were more abundant in organs presenting a large number of MHC-I-peptide complexes. In addition, proteins known to negatively regulate protein degradation were found to be more abundant in organs presenting low numbers of MHC-I peptides. In fact, correlations between protein abundances and numbers of MHC-I peptides detected in tissues were found to be remarkably informative and could be used to systematically infer the role of new proteolytic enzymes in antigen processing. Proteolytic enzymes are critically important in antigen processing. Beside the proteasome, ∼20 proteases act in the MHC-I presentation pathway and can alter presented peptides (Lázaro et al., 2015). ERAP1 is probably the most relevant example here since this aminopeptidase plays a major role in antigen processing through N-terminal peptide trimming into the endoplasmic reticulum and is associated with a number of different autoimmune diseases (Hanson et al., 2018; Serwold et al., 2002). Other aminopeptidases such as leucine aminopeptidase 3 (LAP3) and peptidase D (PEPD) were showcased in this study. Most surprisingly, we identified four carboxypeptidases (CPE, CNDP1, CNDP2 and CPVL)— none of them reported so far to influence the repertoire of MHC-I peptides. These carboxypeptidases might represent new players of the antigen processing and presentation pathway. If tested and validated, such findings would be particularly fascinating because angiotensin-converting enzyme (ACE) is the only ER-resident carboxypeptidase documented so far (Eiseniohr et al., 1992; Shen et al., 2011, 2008), and was shown to be immunologically relevant through production of minor histocompatibility antigens, polyoma virus epitopes and HIV gp160 epitope (Neefjes et al., 2011). The use of chemical inhibitors and CRISPR technology, together with high-throughput immunopeptidomic experiments would be of great value in this context to systematically investigate the role of new proteolytic proteins in shaping the composition and heterogeneity of the MHC-Ii in different cell and tissue types.

Two distinct categories of self-peptides were investigated in this study: those that are tissue-specific and those that are widely presented across most tissues, referred in this study as housekeeping/universal MHC-I peptides. Interestingly, our results show that these two categories of self-peptides share very distinct intrinsic features. The latter is composed of peptides that are highly abundant and strong MHC-I binders in addition to derive from highly expressed genes that are preferentially hyperconserved across evolution. In contrast, tissue-specific peptides are relatively less abundant and are encoded by genes that are strongly expressed in the tissue of origin, but weakly or not expressed in most tissues. Given the distinct properties of those self-peptides, their respective impact toward various immunological processes could be dramatically different, for triggering T cell tolerance in particular. In fact, tolerance mechanisms through recognition of self-peptides, both in the thymus and in the periphery, are critical to eliminate or control self-reactive T cells that would otherwise lead to autoimmunity (Granados et al., 2015; Verteuil et al., 2012; Xing and Hogquist, 2012). Failure to T cell tolerance against housekeeping/universal peptides would have devastating consequences as self-reactive T cells would destroy most organs across the entire organism. Fortunately, we observed that genes coding for those peptides are among the most expressed across entire genomes, hence, increasing the probability that those peptides will be abundantly presented in the thymus to trigger clonal deletion of immature self-reactive T cells recognizing those peptides. Moreover, we made the fundamental observation that housekeeping/universal peptides originate from hyperconserved genes. Therefore, the adaptive immune system may have evolved for 500 million years a remarkable mechanism enabling the elimination of those T cells in a highly efficient manner. In contrast, controlling self-reactivity of T cells recognizing tissue-specific peptides might be more challenging, thereby rationalizing the need for peripheral tolerance processes to avoid tissue-specific autoimmunity (Matsumoto et al., 2019).

Another important observation in this study was that the multiple HLA-I allotypes expressed by a given individual may contribute unevenly to the composition of the MHC-Ii from one organ to another. For instance, HLA-B40:02-associated peptides were found to be particularly enriched in the liver of a given individual compared to all the other organs. Overall, 37 enrichment patterns were observed across 27 specific HLA-I allotypes and 18 different tissues. These are important basic observations because peptide antigens that are processed and presented in a tissue-dependent fashion may cause differential phenotypic consequences in response to the same signal. For instance, in infectious diseases, *Plasmodium* parasites (malaria) and SARS-CoV-2 (COVID-19) have the ability to reach and infect many host tissues (Coban et al., 2018; Wadman et al., 2020). In this context, CD8+ T cells may behave very differently from one tissue to another following tissue-dependent processing and presentation of pathogen-derived peptide antigens, thereby likely impacting the overall efficiency of viral clearance by T cells. Interestingly, tissue-dependent antigen presentation may also lead to a web of tissue-resident memory T cells that functionally adapt to their environment to stop viral spread across the organism (Kadoki et al., 2017). Hence, tissue-specific variations in the MHC-Ii likely play a role in controlling infections or determining the severity of a disease. One can anticipate that immunopeptidomics approaches will be increasingly powerful in the future to investigate the dynamics of the MHC class I antigen processing and presentation pathway *in vivo* and evaluate its impact on tissue-dependent T cell responses in the organism.

Systems understanding of MHC-I antigen presentation at the organism level is at an early stage. MS technologies are constantly evolving and we anticipate that the tissue-specificity of the MHC-Ii will be further refined in the future. In fact, we envision that further improvement in proteomics and immunopeptidomics technologies will enable more robust, precise and comprehensive measurements of proteomes and immunopeptidomes in healthy tissues as well as in response to a wide range of immunological perturbations. Integration of those measurements over time, together with new high-throughput TCR-MHC peptide interaction studies (Dendrou et al., 2018; Moritz et al., 2019), will help understand how widespread and tissue-specific changes in peptide processing and presentation impact tissue-dependent T cell responses, and hence, help understand inter-organ communications between T cell networks to shape the organismal circuitry of immunity (Chevrier, 2019; Kadoki et al., 2017). From a synthetic biology perspective, in-depth understanding of how MHC-I-associated peptides are generated *in vivo* will enable accurate prediction of their dynamics, and ultimately, will accelerate the engineering of new biological systems to control their presentation and function in immunity.

## MATERIALS AND METHODS

### Retrieval and preparation of omics data from the literature

#### Mouse immunopeptidome

Raw data from the mouse immunopeptidome dataset (Schuster et al., 2018) were downloaded and re-analyzed using “PEAKS 9 (Bioinformatics Solutions Inc., Waterloo, Ontario, Canada)” (Tran et al., 2018). Peptides identified with an FDR<5% were further assessed for binding to the MHC-I alleles H2Kb and H2Db using NetMHCpan4.0 (Jurtz et al., 2017). Peptides with a length of 8,9,10,11 or 12 amino acids and a NetMHCpan-4.0 Rank score smaller than 2.0 (Rank <= 2.0) were selected as MHC-I peptides. A collection of all mouse MHC-I peptides is made available in Supplementary Table 1. All downstream data analysis is based on this set of MHC-I peptides.

#### Mouse RNAseq data

Mouse RNAseq data were obtained from (Söllner et al., 2017) supplementary materials and can be found in Supplementary Table 1. Data were used for further analysis in the form provided.

#### Mouse proteomics data

Mouse proteomic data were downloaded from (Geiger et al., 2013) supplementary materials and can be found in Supplementary Table 3. Protein intensities presented in Supplementary Table 3 are the reported label free intensities normalized across tissues.

#### Human immunopeptidome

Human immunopeptidome data were obtained from Marcu et al.,2021 and represent the 2020.06 release of the dataset as can be accessed at https://hla-ligand-atlas.org/. Peptides from this dataset were predicted for HLAI binding using NetMHCpan4.0. Only alleles present in the donor database [based on allele genotyping as described in Marcu et al.] were predicted. Out of six alleles genotyped to each donor, the allele with the lowest NetMHCpan-4.0 Rank score was assigned to a given peptide. For the analysis presented in this manuscript, peptides with a NetMHCpan-4.0 rank smaller or equal than 2 (Rank<=2) were considered MHC-I peptides. The quantitative information, as reported by MHCquant (Bichmann et al., 2019), was also used in the current manuscript. Raw peptide intensities were used as approximative quantitative information and no normalization was performed due to the heterogeneous nature of pulldowns and primary tissue samples. A complete list of peptides including metadata can be found in Supplementary Table 2.

### Human RNAseq data

Human RNAseq data were obtained from the GTEX repository https://www.gtexportal.org/home/ (accessed January 10^th^ 2020), the dataset used was: ‘GTEx_Analysis_2017-06-05_v8_RNASeQCv1.1.9_gene_median_tpm.gct’. The subset of data used for this manuscript can be found in Supplementary Table 2. Data were used for further analysis in the form provided.

### Human proteomics data

Human proteomics data were obtained from (Wang et al., 2019). The subset of data used for this publication can be found in Supplementary Table 4. Data were used for further analysis in the form provided, unless stated differently.

### Principal component analysis

Principal component analysis and visualization was performed in R using the FactoMineR package (Lê et al., 2008). Input variables consist of 19 mouse tissues and 28 human tissues for which immunopeptidomic data are available (note that human thymus tissues were sampled differently and were excluded from this analysis). For each tissue, a vector of individual peptide intensities (log10 transformed) was loaded. The first two dimensions accounting for most of the variability in the data were plotted (Figure 2 D and E).

### Tissue Connectivity Maps

For every possible pair of tissues, the number of overlapping peptides was determined for the mouse and human immunopeptidomes, respectively. A peptide was considered overlapping if an intensity value had been reported in both tissues. A connectivity matrix was generated from the resulting data for mouse and human, respectively (Figure 2F and Supplementary Figure 1). Noteworthily, the number of peptides unique to a given tissue is depicted along the diagonal of depicted heatmaps.

### Tissue-dependent representation of HLA alleles

The proportion of peptides represented by a specific allele in a given tissue was calculated for every subject. Similarly, the mean proportion of every allele across tissues was calculated for every subject. These values were then used to calculate the over- or under-representation of each allele in a tissue compared to the mean as follows:

Subject dependent allele enrichment in tissues:

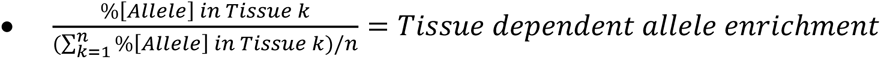

Examples for subject specific allele representations can be found in Figure 2 B-D. In order to assess trends across all subjects, we calculated the mean of these over- and under-representation values for all alleles across all subjects. To find trends among the data, we focused only on alleles over-represented by, on average, at least 1.5-fold in a given tissue across all subjects. Results are depicted in Figure 2 E.

### Connecting mouse immunopeptidomic data with mouse RNAseq data

Source genes of mouse MHC-I peptides available from the Peaks results were mapped to ENSEMBL identifiers using the mouse annotation package org.Mm.eg.db in R (DOI: 10.18129/B9.bioc.org.Mm.eg.db). These source genes were then mapped to the genes in the RNAseq dataset (Söllner et al., 2017) to assess their tissue-dependent RNAseq expression (Supplementary Table 1). All mappings between different gene identifiers were performed using the R package AnnotationHub (DOI: 10.18129/B9.bioc.AnnotationHub).

### Source genes from tissue-specific MHC-I peptides in mouse (Mouse source genes)

Genes mapped to a peptide which is present in only one of the nineteen tissues analyzed in the mouse immunopeptidome are considered to be source genes of tissue specific MHC-I peptides. We have not assessed to what extend additional MHC-I peptides from such a gene are represented across tissues (for genes where more than one MHC-I peptide was identified). We found 2448 source genes from tissue-specific MHC-I peptides in mouse.

### Tissue dependent expression of mouse source genes

In order to visualize the expression of source genes originating from tissue-specific MHC-I peptides, we mapped the 2448 source genes we found in mouse to the mRNA expression atlas published by (Söllner et al., 2017). For the 9 tissues where transcriptomics and proteomics data were available, we extracted mRNA expression levels of the mouse source genes. Expression levels were then averaged across each tissue, grouped by the source tissue (Tissue in which gene represents immunopeptides). Within each source tissue group, a z-score for the average expression value of genes was calculated. The resulting matrix is visualized in Figure 4C.

Note: Z-scores were calculated as follows:

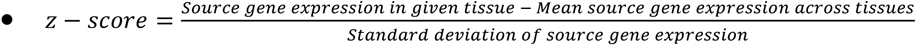

### Conservation of source genes from universal MHC-I peptides (Mouse)

Universal MHC-I peptides are defined as MHC-I peptides present in all of the 19 sampled mouse tissues (Supplementary Figure 6A). Peptides from 38 genes were found in the mouse dataset (Supplementary Table 1). We calculated the conservation of the exon and promoter regions of the corresponding source genes and compared their genetic conservation to those from source genes of tissue specific MHC-I peptides. Conservation scores were extracted in form of PhastCons conservation probabilities (Siepel et al., 2005; Siepel and Haussler, 2004) from publicly available multiple alignments of the mouse genome and the genomes of 59 vertebrates (http://hgdownload.soe.ucsc.edu/goldenPath/mm10/multiz60way/) from the UCSC genome browser (https://genome.ucsc.edu/index.html) (accessed June 5^th^ 2020). BigWig files containing PhastCons scores for the mouse genome were downloaded and queried for the genes of interest using the R package rtracklayer (Lawrence et al., 2009) together with gene positional information from the ‘TxDb.Mmusculus.UCSC.mm10.knownGene’ database provided by the UCSC genome browser (DOI: 10.18129/B9.bioc.TxDb.Mmusculus.UCSC.mm10.knownGene). PhastCons scores for nucleotides of the exon and promoter regions of housekeeping and tissue-specific source genes were extracted from the BigWig files. Promoter regions were defined as 200 bases downstream and 2000 bases upstream of the transcription start site. Conservation scores were then calculated using a 12-base pair sliding window along the extracted genetic regions and the maximum PhastCons value was used as the conservation score. The cumulative frequency of these PhastCons values for the exon and promoter regions of source genes from universal MHC-I peptides and source genes from tissue-specific MHC-I peptides were calculated and compared using the Wilcoxon rank sum test, respectively. This analysis and workflow were inspired by Zeng et al (Zeng et al., 2016) and Zhu et al (Zhu et al., 2008) who investigated the genomic conservation of housekeeping genes compared to tissue specific genes in mouse and human, respectively. Furthermore, ideas for the implication of PhastCons conservation rates were derived from Sun et al. (Sun et al., 2014).

### Annotating the molecular weight of MHC-I peptide source genes (Mouse)

Molecular weights of proteins were retrieved from www.uniprot.org (Complete Mus musculus proteome, reviewed + un-reviewed proteins, accessed June 17 2020). Uniprot identifiers were matched to ENSEMBL gene identifiers and used for analysis.

### Connecting human immunopeptidomic data with human RNAseq data

Source genes of human MHC-I peptides were mapped to ENSEMBL identifiers using the human annotation package org.Hs.eg.db in R (org.Hs.eg.db: Genome wide annotation for Human. R package version 3.8.2). These source genes were then mapped to the genes in the RNAseq dataset (‘GTEx_Analysis_2017-06-05_v8_RNASeQCv1.1.9_gene_median_tpm.gct’) to assess their tissue-dependent RNAseq expression (Supplementary Table 2). All mappings between different gene identifiers were performed using the R package AnnotationHub (DOI: 10.18129/B9.bioc.AnnotationHub).

### Source genes from tissue-specific MHC-I peptides (Human)

Source genes representing one or more MHC-I peptides that were measured in only one tissue sample in the human immunopeptidome dataset were considered source genes from tissue-specific MHC-I peptides. In human we found 12,095 of such genes. Similar to the mouse analysis, we did not assess to what extendt these genes yield additional peptides present in more than one tissue sample.

### Conservation of source genes from universal MHC-I peptides (Human)

Defining source genes from universal MHC-I peptides in human is less straightforward compared to the mouse due to the heterogeneity of subjects from which tissues were sampled and HLA alleles representation. Hence, we defined a source gene from universal MHC-I peptides in the available human immunopeptidome as a gene for which one or more MHC-I peptides were either 1) present across all tissues in at least two patients or 2) present across all samples in which the assigned HLA allele was present or 3) among the top 100 peptides identified the most frequently across all measured samples, independent of allele or subject (Supplementary Figure 6B-F). In order to avoid a bias towars peptides from donors where only few tissues were sampled, we focused only on donors where 14 or more tissues were sampled. This analysis resulted in a total of 251 source genes from universal MHC-I peptides (Supplementary Figure 7 and Supplementary Table 2).

Conservation analysis was performed using PhastCons retrieved from an alignment of the hg38 human genome with 99 vertebrates. Data were downloaded from the UCSC genome browser at http://hgdownload.soe.ucsc.edu/goldenPath/hg38/multiz100way/ (accessed June 5^th^ 2020). Genetic positions of genes of interest (genes from universal and tissue-specific MHC-I peptides) were mapped using the ‘TxDb.Hsapiens.UCSC.hg38.knownGene’ database (DOI: 10.18129/B9.bioc.TxDb.Hsapiens.UCSC.hg38.knownGene) and conservation scores were calculated and compared the same way as the mouse conservation scores.

### Annotating the molecular weight of MHC-I peptide source genes (Human)

Molecular weights of proteins were retrieved from www.uniprot.org (Complete Homo sapiens proteome, reviewed + un-reviewed proteins, accessed June 17 2020). Uniprot identifiers were matched to ENSEMBL gene identifiers and used for analysis.

### Computing and analyzing protein wise correlation between tissue MHC-I peptide counts and protein abundances in mouse and human

Protein expression data from mouse and human proteomic tissue drafts (Supplementary Tables 3 and 4) were obtained from (Geiger et al., 2013) for mouse and (Wang et al., 2019) for human. Both datasets were chosen due to their recency and wide range of tissues sampled. Correlations between the expression pattern of a given protein across tissues and the overall number of MHC-I peptides sampled across tissues in mouse or human subjects were measured. Expression values (log10 transformed) of each protein across tissues were plotted against the number of total MHC-I peptides identified in each tissue and R-squared and p-values were computed if more than 9 measurement pairs (expression value and total number of MHC-I peptides) were available. For the human dataset, correlations were calculated for immunopeptidome data from every subject where above criteria were fulfilled. Expression values from jejunum and duodenum from the human proteomics dataset (Wang et al., 2019) were averaged and paired with total MHC-I peptide counts in the small intestine. P-values and R-squared values were reported. For the human data, we required p-values < 0.05 and R-squared values >0.4 in at least two patients to consider a correlation to be non-random (Supplementary Figure 9A). For the mouse data where only one donor is available, correlations with p-values < 0.01 and R-squared > 0.4 were considered to be non-random observations (Supplementary Figure 9B). Correlation data for all proteins of the mouse and human datasets can be found in Supplementary Tables 3 and 4, respectively.

### Functional proteomic analysis

Gene set enrichment analysis (GSEA; http://www.broad.mit.edu/gsea/) was performed using GSEA software and the Molecular Signature Database (MsigDB) on proteins from systematic cross-tissue analysis of MHC class I peptides and protein expression. Top 50 significant gene sets using the Gene Ontology modules overlap analysis were considered significant with P value and FDR < 0.05. We acknowledge our use of the GSEA, GSEA software, and MSigDB (Subramanian et al., 2005). Results can be found in Supplementary Table 5.

### Code availability in form of the MHCIatlas R package

In order to achieve reproducibility of the presented data analysis, to allow further and flexible data analysis by readers and to make the code used publicly available, we generated an R-package entitled ‘MHCIatlas’ accompanying this manuscript. The package can be accessed from the GitHub page of the ‘CaronLab’ at https://github.com/CaronLab/MHCIatlas. Download of the source code can be performed directly from the above link. Alternatively, the package can be installed in R using the ‘install_github’ function from the ‘devtools’ package as shown below:

~~~
> devtools::install_github(‘CaronLab/MHCIatlas’)
~~~

The ‘MHCIatlas’ R package includes 30 functions to reproduce the data-analysis presented in this manuscript as well as the immunopeptidomic, proteomics and transcriptomics datasets used. A quick guide to install and use the R package ‘MHCIatlas’ is provided as supplementary material in form of the ‘MHCIatlas user guide’ pdf document.

## ACKNOWLEDGMENT

This study was supported by funding from the Fonds de recherche du Québec – Santé (FRQS), the Cole Foundation, CHU Sainte-Justine and the Charles-Bruneau Foundations, Canada Foundation for Innovation and by the National Sciences and Engineering Research Council (NSERC) (#RGPIN-2020-05232). This work was also funded by the Deutsche Forschungsgemeinschaft (DFG, German Research Foundation) under Germany’s Excellence Strategy - EXC 2180 – 390900677; the Deutsche Forschungsgemeinschaft (DFG) SFB 685 „Immunotherapy: Molecular Basis and Clinical Application“; the ERC AdG 339842 MUTAEDITING; the Boehringer Ingelheim Foundation for Basic Research in Medicine, the Bosch Research Foundation, and the German Network for Bioinformatics Infrastructure (de.NBI). K.K. is a recipient of IVADO’s postdoctoral scholarship (#4879287150). E.C. is a FRQS Junior 1 Research Scholar.

## AUTHOR CONTRIBUTIONS

PK and IS analyzed the data. AM, LB, LK and HS generated and analyzed the data. EC and PK wrote the manuscript and all the co-authors provided critical comments.

## SUPPLEMENTARY FIGURES

**Supplementary Figure: 1:**
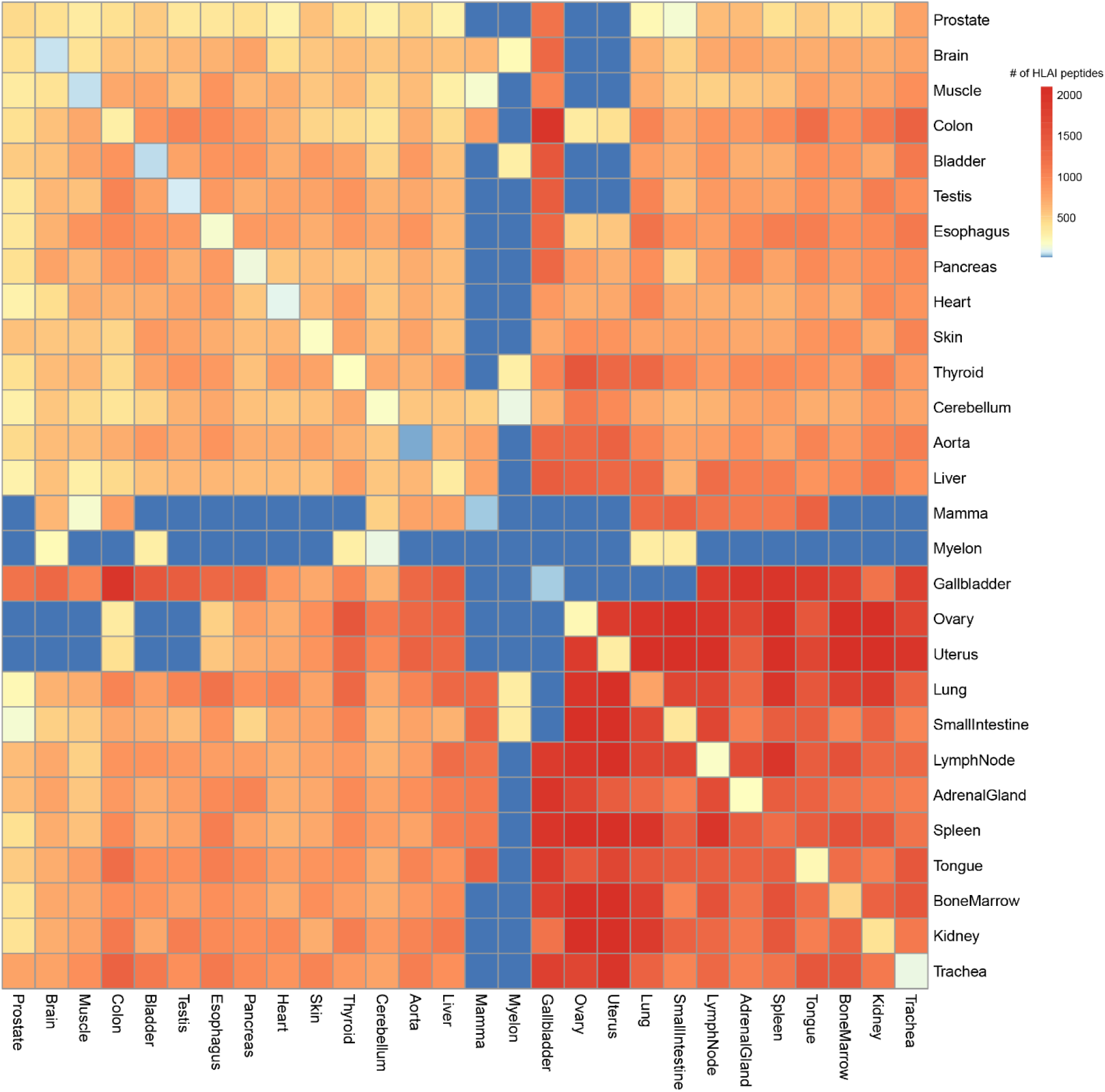
Connectivity map human immunopeptidome. This human heat map integrates immunopeptidomic data from 38 HLAI allotypes and 13 different subjects. The human heat map is not deconvoluted by HLA allotype nor subject and therefore provide a bird’s eye view of the human class I immunopeptidome. Note: The number of uniquely observed/tissue-specific peptides can be found along the diagonal.

**Supplementary Figure 2:**
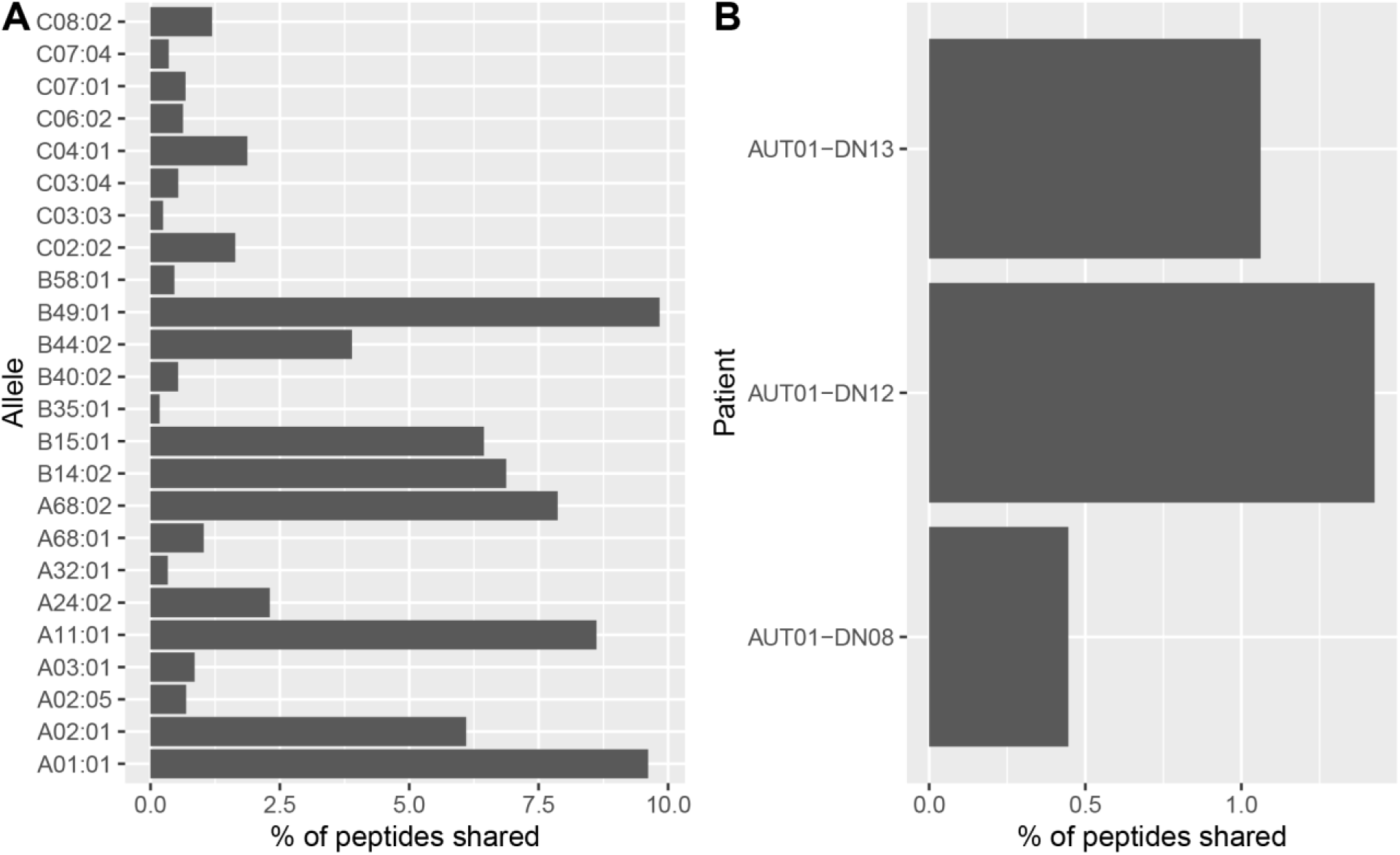
Proportion of peptides shared across Colon, Spleen, Liver, Lung, Bone marrow and Kidney. **(A)** Deconvoluted by best allele for which all 6 tissues were sampled. **(B)** Deconvoluted by subjects for which all six tissues were sampled.

**Supplementary Figure 3:**
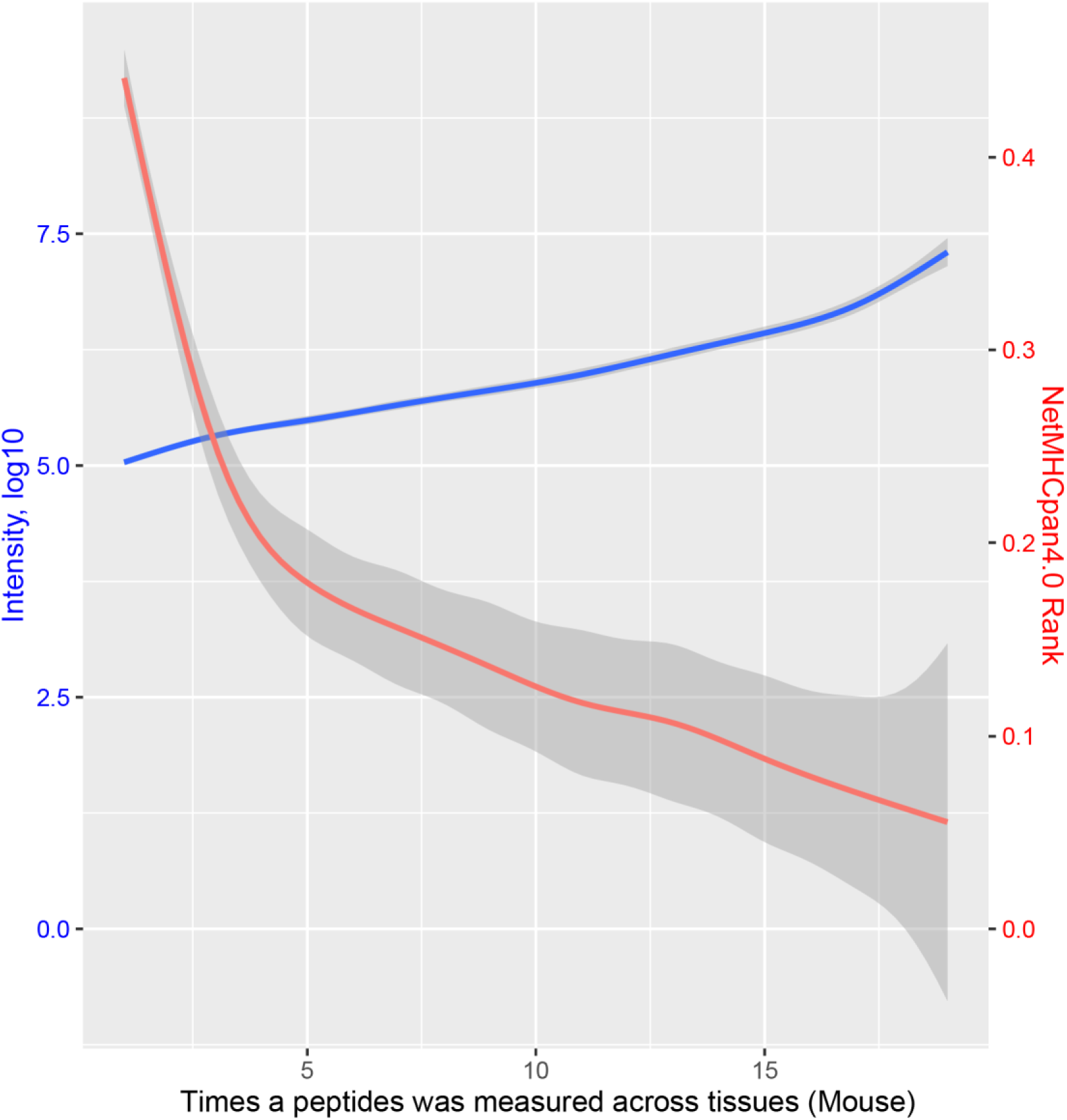
NetMHCpan4.0 rank and MHCI peptide intensities plotted against the number of measurements across tissues in the mouse dataset.

**Supplementary Figure 4:**
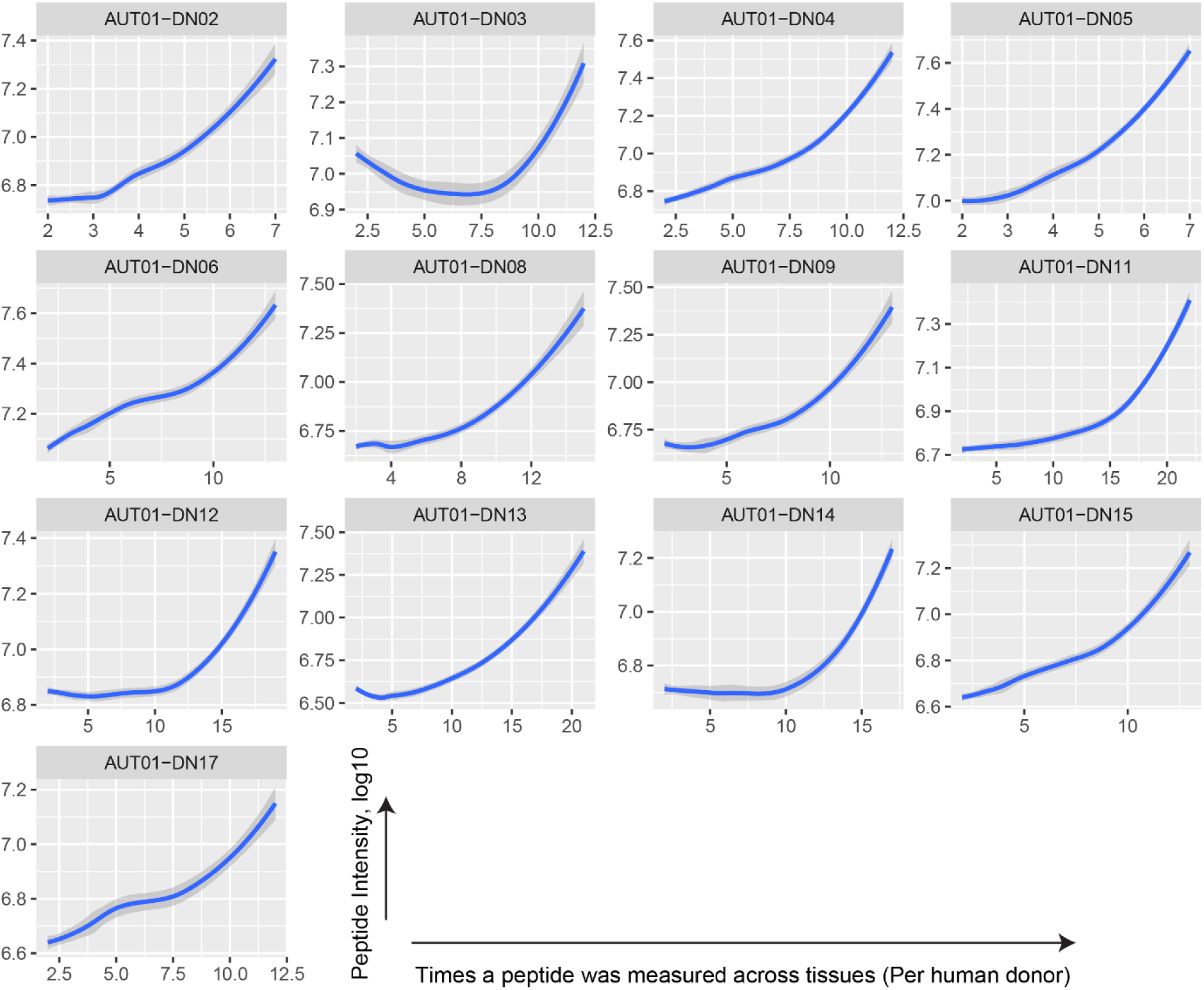
MHC I peptide intensities plotted against the number of measurements across tissues in the human dataset. Each panel represents one subject.

**Supplementary Figure 5:**
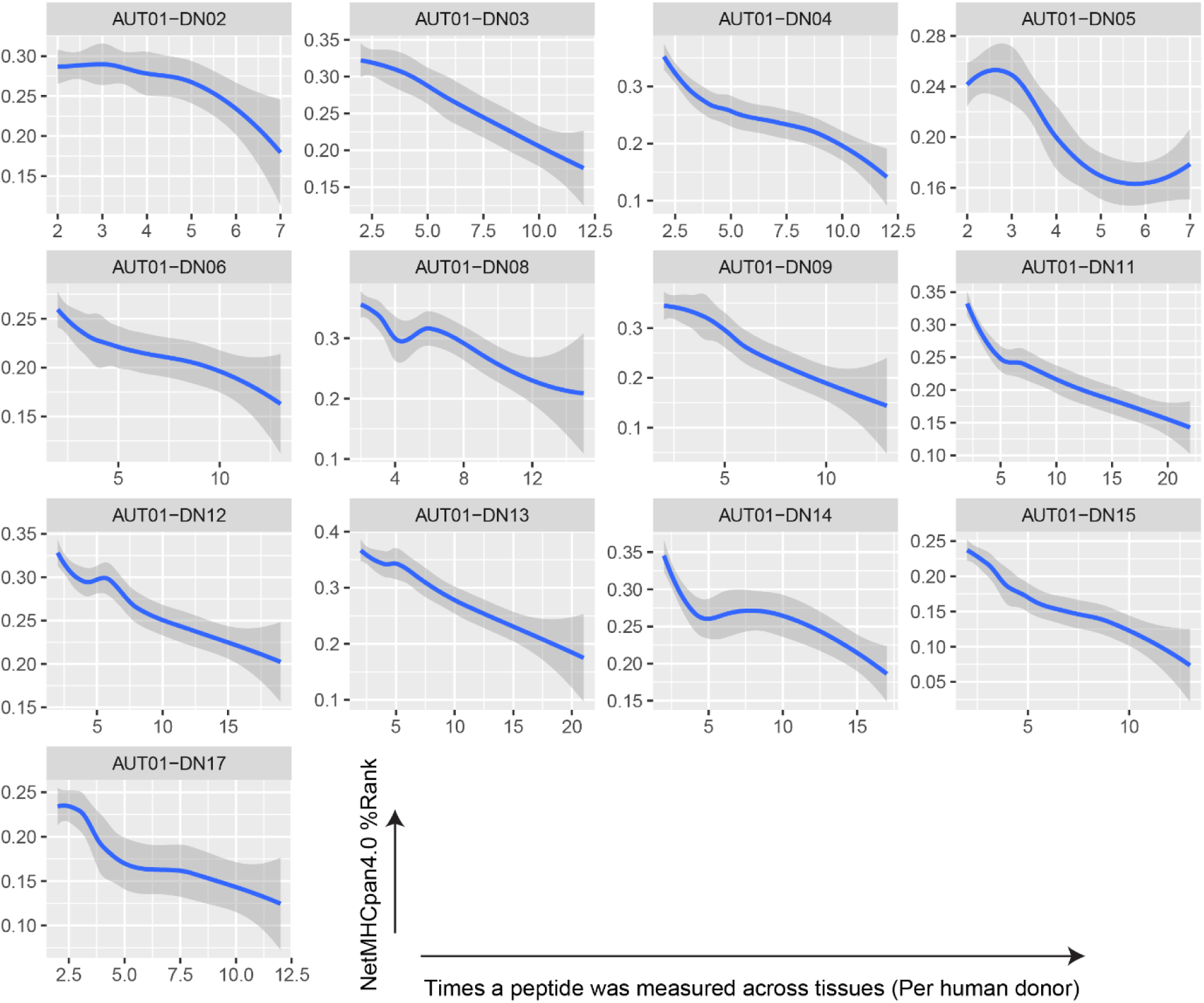
NetMHCpan4.0 rank (best allele) plotted against the number of measurements across tissues in the human dataset. Each panel represents one subject.

**Supplementary Figure 6:**
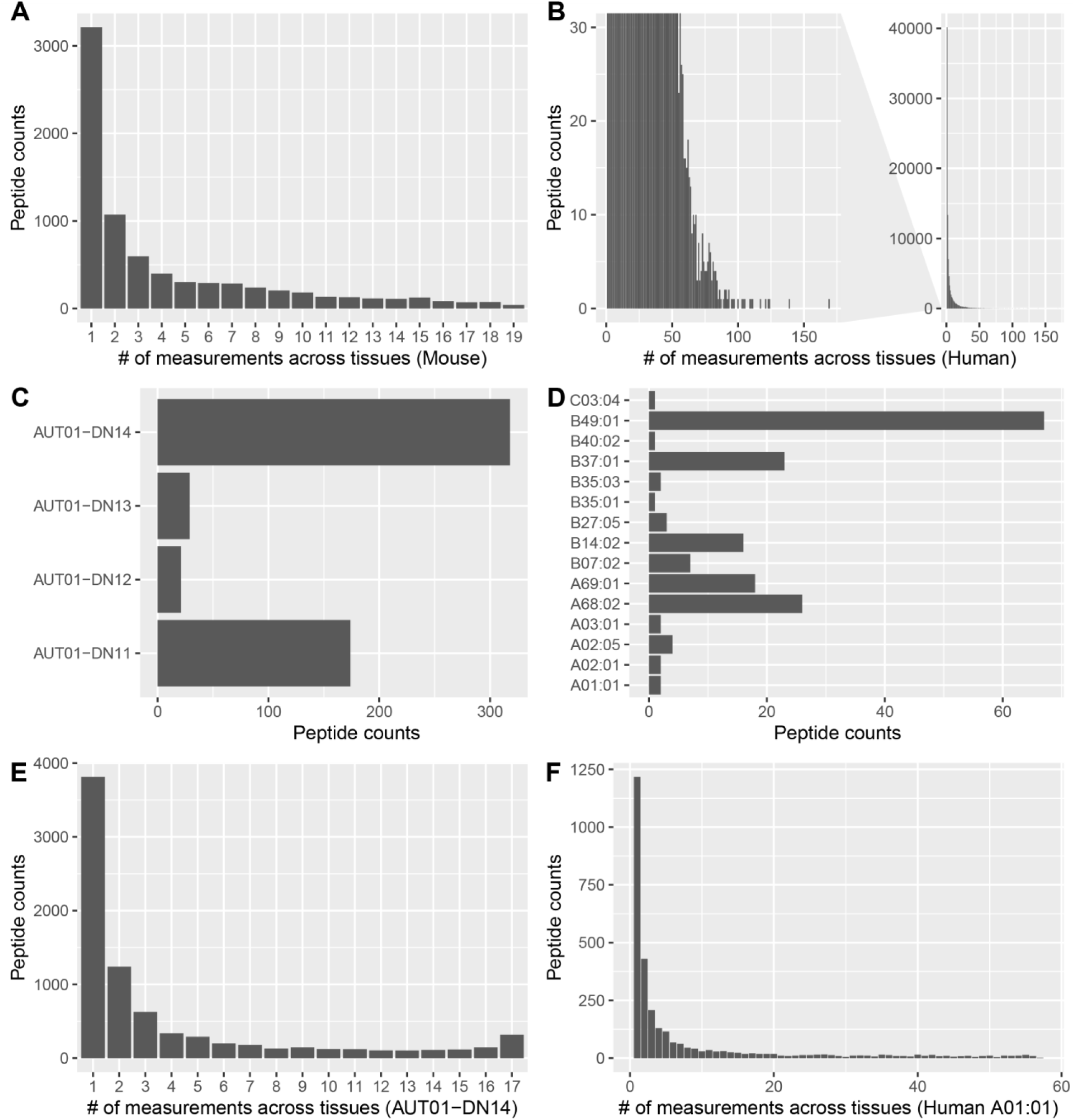
Identification of the mouse and human tissue-independent MHCI peptides and their corresponding source genes. **(A)** Histogram showing the number of MHCI peptides presented in one or more tissues. 1 measurement are MHCI peptides identified in only one tissue whereas 19 measurements are MHCI peptides identified in all the 19 tissues. Mouse source genes coding for MHCI peptides that were measured across more than 17 tissues were considered for further analysis. **(B)** Source genes of the top 100 peptides from the human dataset measured the most frequently across tissues were considered ‘source genes of universal peptides’. **(C)** In addition to B, source genes that present peptides across all tissues in at least two patients are also considered as ‘source genes of universal peptides’. **(D)** In addition to B and C, source genes that present peptides across all tissues for at least one allele are considered ‘source genes of universal peptides’. **(E)** Example of the distribution of peptide counts across tissues in AUT-DN14. **(F)** Example distribution of peptide counts across all samples containing allele A01:01.

**Supplementary Figure 7:**
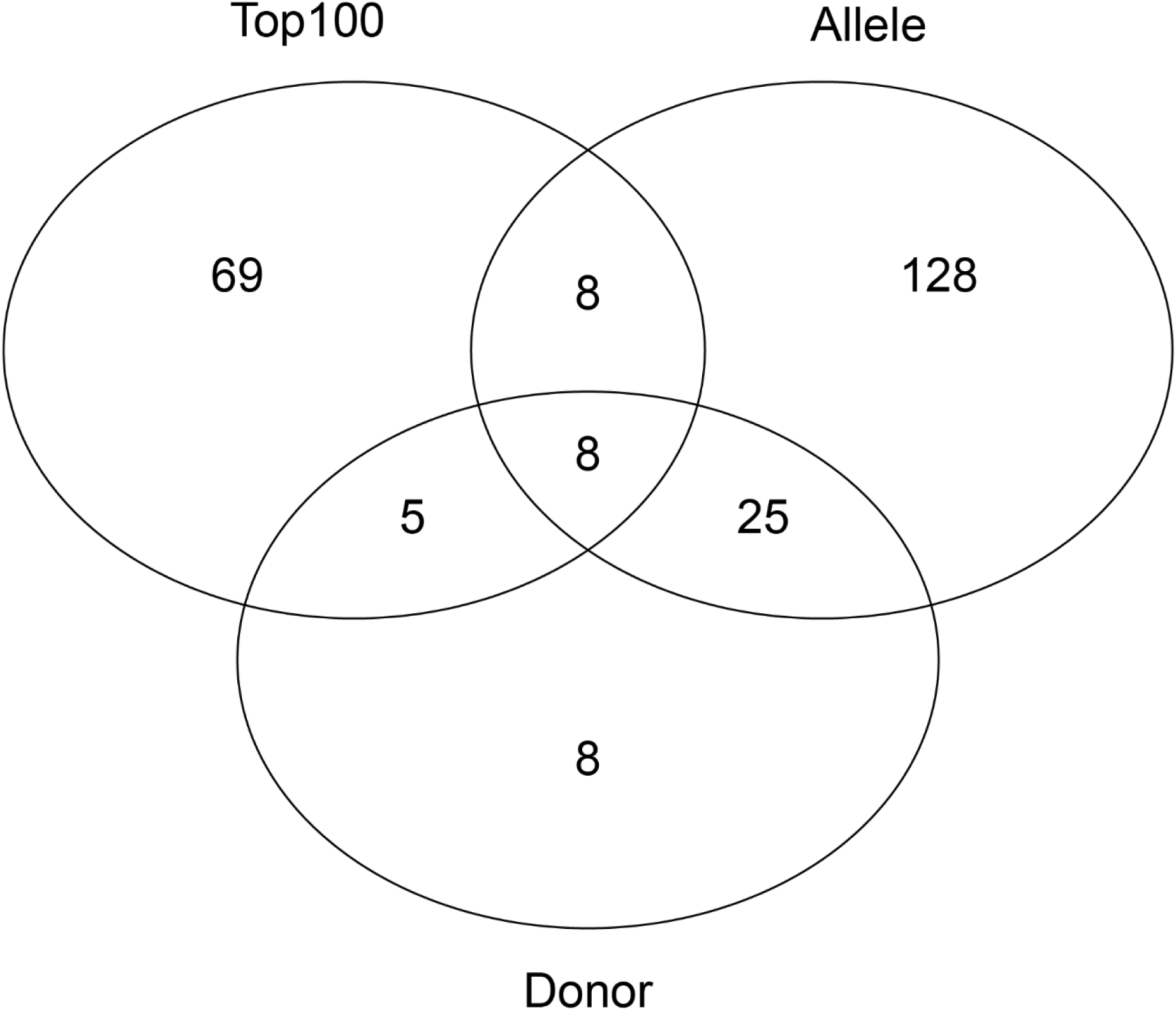
Venn diagram of human universal-peptide source genes originating from the three selection criteria specified in Supplementary Figure 6 B-D.

**Supplementary Figure 8:**
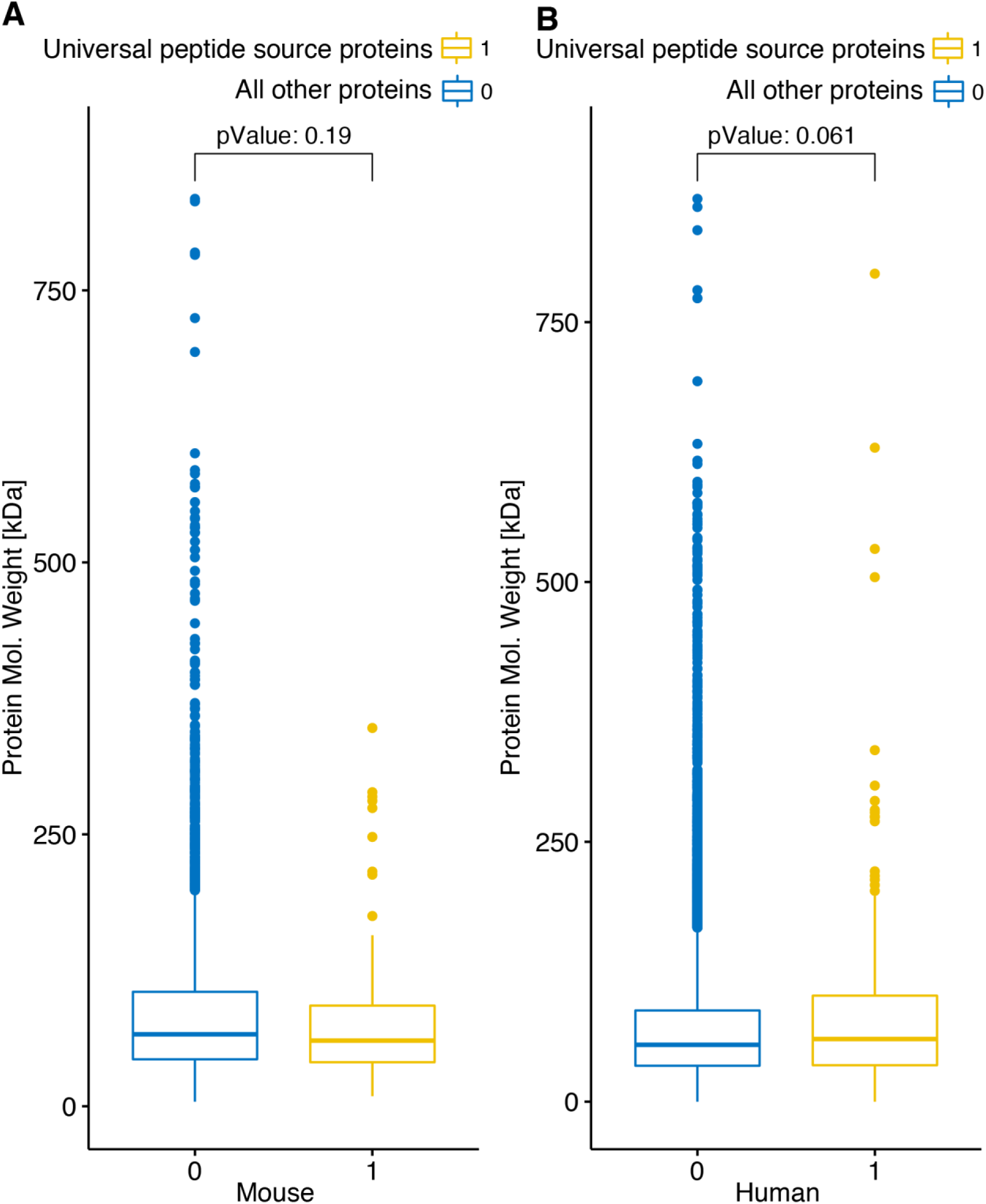
Molecular weight of universal-peptide and tissue-specific-peptide source proteins. **(A)** Mouse and **(B)** Human.

**Supplementary Figure 9:**
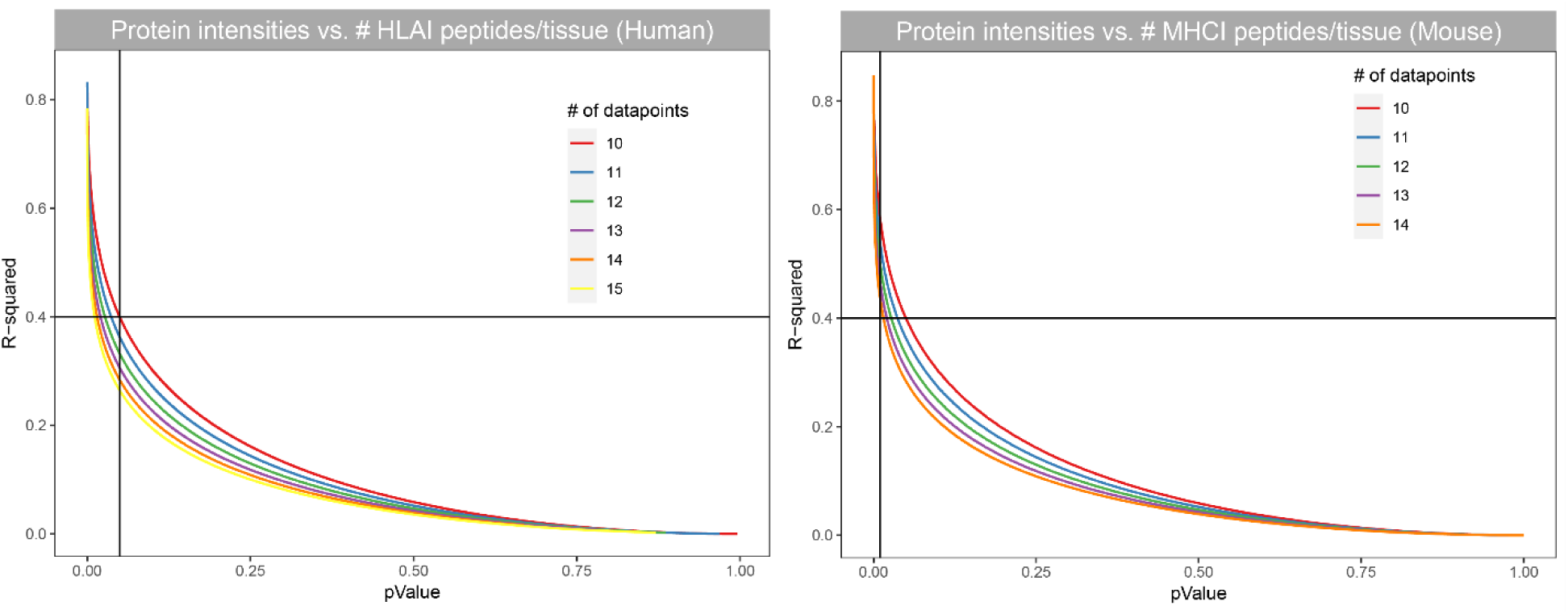
Large scale correlation of protein intensities with the total count of MHCI peptides per tissue in human and mouse datasets. **(A)** R-squared of linear fits plotted against the corresponding p-values for the human data. Proteins whose fits show R-squared values> 0.4 (p-value<0.05) in at least two subjects are considered significant. **(B)** R-squared of linear fits plotted against the corresponding p-values for the mouse data. Proteins whose fits show R-squared values> 0.4 (p-value<0.01) are considered significant.

**Supplementary Figure 10:**
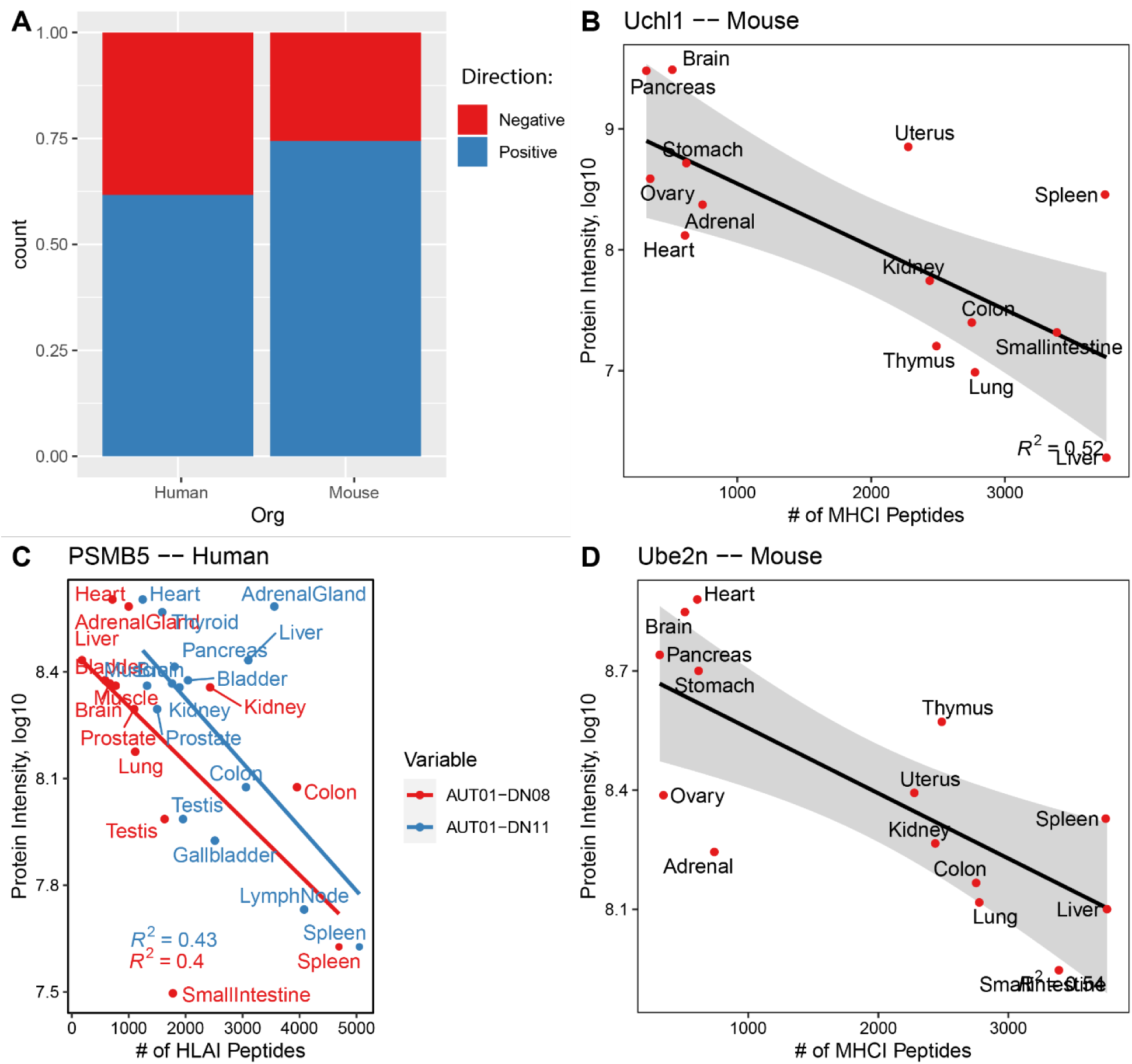
Negative correlations between total MHC-I peptide counts and protein intensities. **(A)** Proportion of direction of slopes of significant proteins in human and mouse. **(B-D)** Example fits of proteins in human and mouse with negative correlation.

**Supplementary Figure 11:**
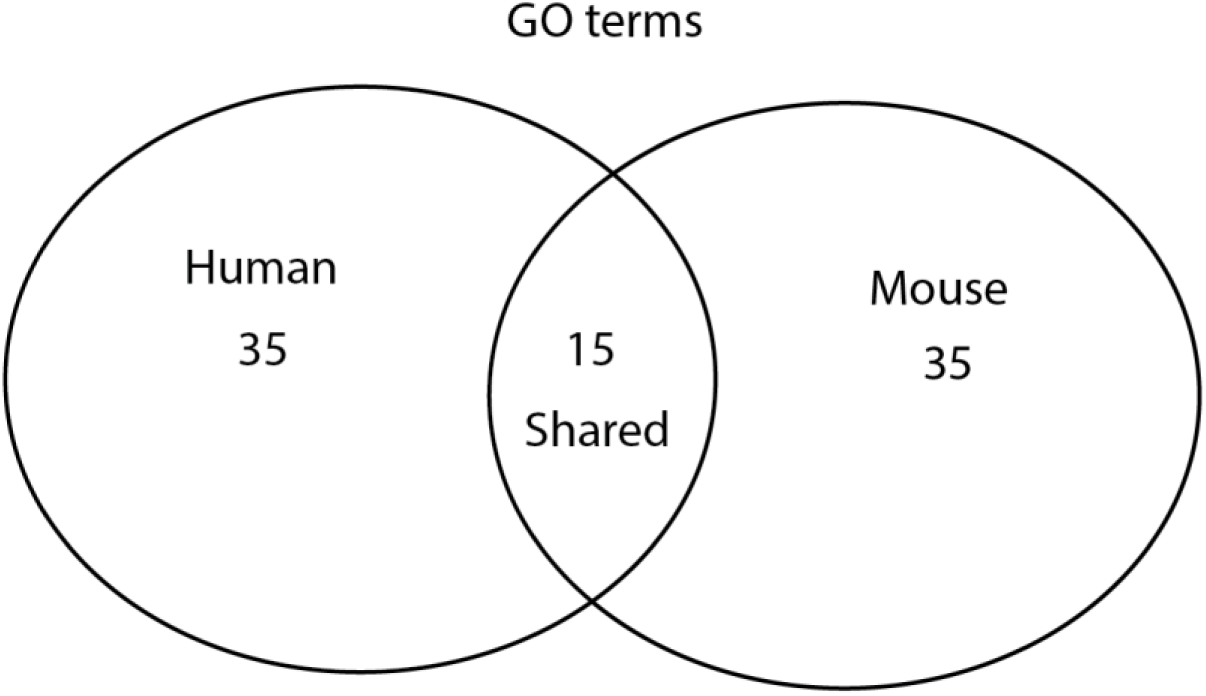
Overlap of enriched gene ontology (GO) terms between Mouse and Human for genes significantly correlating with total MHCI/HLAI counts.

**Supplementary Figure 12:**
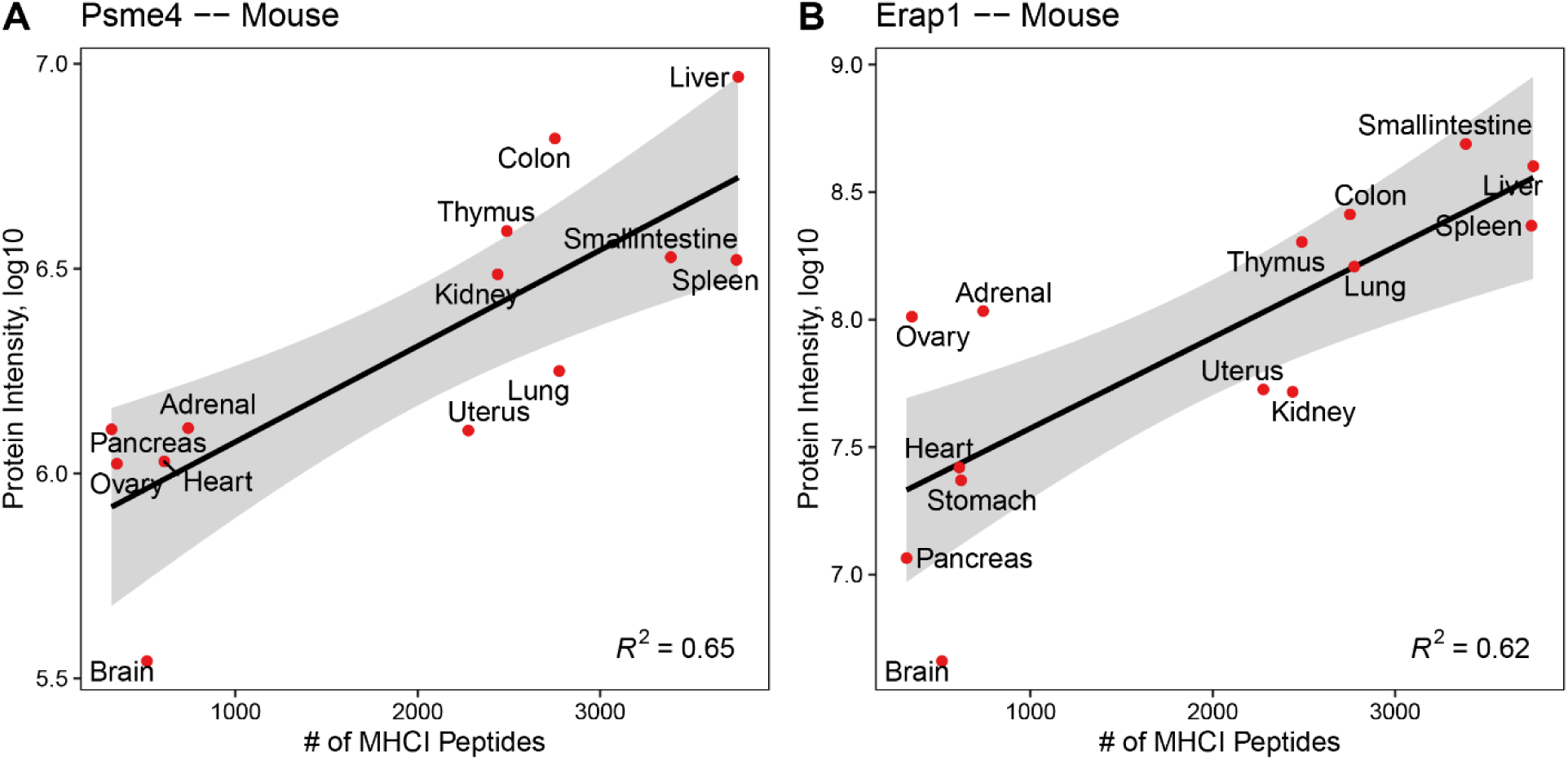
Example fit curves of prominent proteins. **(A)** Example fit of the protein Psme4 in mouse. **(B)** Example fit of the protein Erap1 in mouse.

## REFERENCES

Abelin JG, Keskin DB, Sarkizova S, Hartigan CR, Zhang W, Sidney J, Stevens J, Lane W, Zhang GL, Eisenhaure TM, Clauser KR, Hacohen N, Rooney MS, Carr SA, Wu CJ. 2017. Mass Spectrometry Profiling of HLA-Associated Peptidomes in Mono-allelic Cells Enables More Accurate Epitope Prediction. Immunity 46:315–326. doi:10.1016/j.immuni.2017.02.007

Alvarez-Navarro C, Martin-Esteban A, Barnea E. 2015. ERAP1 polymorphism relevant to inflammatory disease shapes the peptidome of the birdshot chorioretinopathy-associated HLA-A*29:02 antigen. Mol Cell Proteomics 14:1770–1780. doi: 10.1074/mcp.M115.048959.

Bassani-Sternberg M, Pletscher-Frankild S, Jensen LJ, Mann M. 2015. Mass Spectrometry of Human Leukocyte Antigen Class I Peptidomes Reveals Strong Effects of Protein Abundance and Turnover on Antigen Presentation. Mol Cell Proteomics 14:658–673. doi:10.1074/mcp.m114.042812

Bichmann L, Nelde A, Ghosh M, Heumos L, Mohr C, Peltzer A, Kuchenbecker L, Sachsenberg T, Walz JS, Stevanović S, Rammensee H-G, Kohlbacher O. 2019. MHCquant: Automated and reproducible data analysis for immunopeptidomics. J Proteome Res. 18:3876–3884. doi:10.1021/acs.jproteome.9b00313

Blees A, Januliene D, Hofmann T, Koller N, Schmidt C, Trowitzsch S, Moeller A, Tampé R.2017. Structure of the human MHC-I peptide-loading complex. Nature 551:525–528. doi:10.1038/nature24627

Bourdetsky D, Schmelzer CEH, Admon A. 2014. The nature and extent of contributions by defective ribosome products to the HLA peptidome. Proc Natl Acad Sci USA 111:E1591–E1599. doi:10.1073/pnas.1321902111

Burr ML, Sparbier CE, Chan KL, Chan Y-C, Kersbergen A, Lam EYN, Azidis-Yates E, Vassiliadis D, Bell CC, Gilan O, Jackson S, Tan L, Wong SQ, Hollizeck S, Michalak EM, Siddle HV, McCabe MT, Prinjha RK, Guerra GR, Solomon BJ, Sandhu S, Dawson S-J, Beavis PA, Tothill RW, Cullinane C, Lehner PJ, Sutherland KD, Dawson MA. 2019. An Evolutionarily Conserved Function of Polycomb Silences the MHC Class I Antigen Presentation Pathway and Enables Immune Evasion in Cancer. Cancer Cell 36:385–401.e8. doi:10.1016/j.ccell.2019.08.008

Caron E, Aebersold R, Banaei-Esfahani A, Chong C, Bassani-Sternberg M. 2017. A Case for a Human Immuno-Peptidome Project Consortium. Immunity 47:203–208. doi:10.1016/j.immuni.2017.07.010

Caron E, Espona L, Kowalewski DJ, Schuster H, Ternette N, Alpízar A, Schittenhelm RB, Ramarathinam SH, Arlehamn CSL, Koh CC, Gillet LC, Rabsteyn A, Navarro P, Kim S, Lam H, Sturm T, Marcilla M, Sette A, Campbell DS, Deutsch EW, Moritz RL, Purcell AW, Rammensee H-G, Stevanovic S, Aebersold R. 2015a. An open-source computational and data resource to analyze digital maps of immunopeptidomes. Elife 4. doi:10.7554/elife.07661

Caron E, Kowalewski DJ, Koh CC, Sturm T, Schuster H, Aebersold R. 2015b. Analysis of Major Histocompatibility Complex (MHC) Immunopeptidomes Using Mass Spectrometry. Mol Cell Proteomics 14:3105–3117. doi:10.1074/mcp.o115.052431

Chevrier N. 2019. Decoding the Body Language of Immunity: Tackling the Immune System at the Organism Level. Curr Opin Syst Biology 18:19–26. doi:10.1016/j.coisb.2019.10.010

Coban C, Lee MSJ, Ishii KJ. 2018. Tissue-specific immunopathology during malaria infection.Nat Rev Immunol 18:266–278. doi:10.1038/nri.2017.138

Dendrou CA, Petersen J, Rossjohn J, Fugger L. 2018. HLA variation and disease. Nat Rev Immunol 18:325–339. doi:10.1038/nri.2017.143

Eiseniohr LC, Bacik I, Bennink JR, Bernstein K, Yewdell JW. 1992. Expression of a membrane protease enhances presentation of endogenous antigens to MHC class I-restricted T lymphocytes. Cell 71:963–972. doi:10.1016/0092-8674(92)90392-p

Elenich LA, Nandi D, Kent AE, McCluskey TS, Cruz M, Iyer MN, Woodward EC, Conn CW, Ochoa AL, Ginsburg DB, Monaco JJ. 1999. The complete primary structure of mouse 20S proteasomes. Immunogenetics 49:835–842. doi:10.1007/s002510050562

Falk K, Rötzschke O, Stevanovic S, Jung G, Rammensee H-G. 1991. Allele-specific motifs revealed by sequencing of self-peptides eluted from MHC molecules. Nature 351:290–296. doi:10.1038/351290a0

Fortier M-H, Caron É, Hardy M-P, Voisin G, Lemieux S, Perreault C, Thibault P. 2008. The MHC class I peptide repertoire is molded by the transcriptome. J Exp Med 205:595–610. doi:10.1084/jem.20071985

Geiger T, Velic A, Macek B, Lundberg E, Kampf C, Nagaraj N, Uhlen M, Cox J, Mann M. 2013. Initial Quantitative Proteomic Map of 28 Mouse Tissues Using the SILAC Mouse. Mol Cell Proteomics 12:1709–1722. doi:10.1074/mcp.m112.024919

Gfeller D, Bassani-Sternberg M. 2018. Predicting Antigen Presentation-What Could We Learn From a Million Peptides? Front Immunol 9:1716. doi:10.3389/fimmu.2018.01716

Granados DP, Laumont CM, Thibault P, Perreault C. 2015. The nature of self for T cells—a systems-level perspective. Curr Opin Immunol 34:1–8. doi:10.1016/j.coi.2014.10.012

Granados DP, Yahyaoui W, Laumont CM, Daouda T, Muratore-Schroeder TL, Côté C, Laverdure J-P, Lemieux S, Thibault P, Perreault C. 2012. MHC I–associated peptides preferentially derive from transcripts bearing miRNA response elements. Blood 119:e181–e191. doi:10.1182/blood-2012-02-412593

Hanson AL, Morton CJ, Parker MW, Bessette D, Kenna TJ. 2018. The genetics, structure and function of the M1 aminopeptidase oxytocinase subfamily and their therapeutic potential in immune-mediated disease. Hum Immunol 80:281–289. doi:10.1016/j.humimm.2018.11.002

Huber EM, Basler M, Schwab R, Heinemeyer W, Kirk CJ, Groettrup M, Groll M. 2012.Immuno- and Constitutive Proteasome Crystal Structures Reveal Differences in Substrate and Inhibitor Specificity. Cell 148:727–738. doi:10.1016/j.cell.2011.12.030

Jurtz V, Paul S, Andreatta M, Marcatili P, Peters B, Nielsen M. 2017. NetMHCpan-4.0: Improved Peptide–MHC Class I Interaction Predictions Integrating Eluted Ligand and Peptide Binding Affinity Data. J Immunol 199:3360–3368. doi:10.4049/jimmunol.1700893

Kadoki M, Patil A, Thaiss CC, Brooks DJ, Pandey S, Deep D, Alvarez D, Andrian UH von, Wagers AJ, Nakai K, Mikkelsen TS, Soumillon M, Chevrier N. 2017. Organism-Level Analysis of Vaccination Reveals Networks of Protection across Tissues. Cell 171:398–413.e21. doi:10.1016/j.cell.2017.08.024

Kincaid EZ, Che JW, York I, Escobar H, Reyes-Vargas E, Delgado JC, Welsh RM, Karow ML, Murphy AJ, Valenzuela DM, Yancopoulos GD, Rock KL. 2011. Mice completely lacking immunoproteasomes show major changes in antigen presentation. Nat Immunol 13:129–135. doi:10.1038/ni.2203

Laumont CM, Vincent K, Hesnard L, Audemard É, Bonneil É, Laverdure J-P, Gendron P, Courcelles M, Hardy M-P, Côté C, Durette C, St-Pierre C, Benhammadi M, Lanoix J, Vobecky S, Haddad E, Lemieux S, Thibault P, Perreault C. 2018. Noncoding regions are the main source of targetable tumor-specific antigens. Sci Transl Med 10:eaau5516. doi:10.1126/scitranslmed.aau5516

Lawrence M, Gentleman R, Carey V. 2009. rtracklayer: an R package for interfacing with genome browsers. Bioinformatics 25:1841–1842. doi:10.1093/bioinformatics/btp328

Lázaro S, Gamarra D, Val MD. 2015. Proteolytic enzymes involved in MHC class I antigen processing: A guerrilla army that partners with the proteasome. Mol Immunol 68:72–76. doi:10.1016/j.molimm.2015.04.014

Lê S, Josse J, Husson F. 2008. FactoMineR : An R Package for Multivariate Analysis. J Stat Softw 25. doi:10.18637/jss.v025.i01

Lee CM, Barber GP, Casper J, Clawson H, Diekhans M, Gonzalez JN, Hinrichs AS, Lee BT, Nassar LR, Powell CC, Raney BJ, Rosenbloom KR, Schmelter D, Speir ML, Zweig AS, Haussler D, Haeussler M, Kuhn RM, Kent WJ. 2020. UCSC Genome Browser enters 20th year. Nucleic Acids Res 48:D756–D761. doi:10.1093/nar/gkz1012

Maccari G, Robinson J, Ballingall K, Guethlein LA, Grimholt U, Kaufman J, Ho C-S, de Groot NG, Flicek P, Bontrop RE, Hammond JA, Marsh SGE. 2017. IPD-MHC 2.0: an improved inter-species database for the study of the major histocompatibility complex. Nucleic Acids Res 45:D860–D864. doi:10.1093/nar/gkw1050

Marcu A, Bichmann L, Kuchenbecker L, Kowalewski DJ, Freudenmann LK, Backert L, Mühlenbruch L, Szolek A, Lübke M, Wagner P, Engler T, Matovina S, Wang J, Hauri-Hohl M, Martin R, Kapolou K, Walz JS, Velz J, Moch H, Regli L, Silginer M, Weller M, Löffler MW, Erhard F, Schlosser A, Kohlbacher O, Stevanović S, Rammensee H-G, Neidert MC. 2021. The HLA Ligand Atlas: A benign reference of HLA-presented peptides to improve T-cell-based cancer immunotherapy. J Immunother Cancer, in press

Matsumoto Minoru, Tsuneyama K, Morimoto J, Hosomichi K, Matsumoto Mitsuru, Nishijima H. 2019. Tissue-specific autoimmunity controlled by Aire in thymic and peripheral tolerance mechanism. Int Immunol 32:117–131. doi:10.1093/intimm/dxz066

Milner E, Gutter-Kapon L, Bassani-Strenberg M, Barnea E, Beer I, Admon A. 2013. The effect of proteasome inhibition on the generation of the human leukocyte antigen (HLA) peptidome. Mol Cell Proteomics 12:1853–1864. doi:10.1074/mcp.m112.026013

Moritz A, Anjanappa R, Wagner C, Bunk S, Hofmann M, Pszolla G, Saikia A, Garcia-Alai M, Meijers R, Rammensee H-G, Springer S, Maurer D. 2019. High-throughput peptide-MHC complex generation and kinetic screenings of TCRs with peptide-receptive HLA-A*02:01 molecules. Sci Immunol 4:eaav0860. doi:10.1126/sciimmunol.aav0860

Müller M, Gfeller D, Coukos G, Bassani-Sternberg M. 2017. “Hotspots” of Antigen Presentation Revealed by Human Leukocyte Antigen Ligandomics for Neoantigen Prioritization. Front Immunol 8:1367. doi:10.3389/fimmu.2017.01367

Murata S, Takahama Y, Kasahara M, Tanaka K. 2018. The immunoproteasome and thymoproteasome: functions, evolution and human disease. Nat Immunol 19:923–931. doi:10.1038/s41590-018-0186-z

Nagarajan NA, Verteuil DA de, Sriranganadane D, Yahyaoui W, Thibault P, Perreault C, Shastri N. 2016. ERAAP Shapes the Peptidome Associated with Classical and Nonclassical MHC Class I Molecules. J Immunol 197:1035–1043. doi:10.4049/jimmunol.1500654

Neefjes J, Jongsma MLM, Paul P, Bakke O. 2011. Towards a systems understanding of MHC class I and MHC class II antigen presentation. Nat Rev Immunol 11:823. doi:10.1038/nri3084

Paul P, van den Hoorn T, Jongsma MLM, Bakker MJ, Hengeveld R, Janssen L, Cresswell P, Egan DA, van Ham M, ten Brinke A, Ovaa H, Beijersbergen RL, Kuijl C, Neefjes J. 2011. A Genome-wide Multidimensional RNAi Screen Reveals Pathways Controlling MHC Class II Antigen Presentation. Cell 145:268–283. doi:10.1016/j.cell.2011.03.023

Pearson H, Daouda T, Granados DP, Durette C, Bonneil E, Courcelles M, Rodenbrock A, Laverdure J-P, Côté C, Mader S, Lemieux S, Thibault P, Perreault C. 2016. MHC class I–associated peptides derive from selective regions of the human genome. J Clin Invest 126:4690–4701. doi:10.1172/jci88590

Rêgo AT, Fonseca PCA da. 2019. Characterization of Fully Recombinant Human 20S and 20S-PA200 Proteasome Complexes. Mol Cell 76:138–147.e5. doi:10.1016/j.molcel.2019.07.014

Reits E, Neijssen J, Herberts C, Benckhuijsen W, Janssen L, Drijfhout JW, Neefjes J. 2004. A Major Role for TPPII in Trimming Proteasomal Degradation Products for MHC Class I Antigen Presentation. Immunity 20:495–506. doi:10.1016/s1074-7613(04)00074-3

Reits EAJ, Vos JC, Grommé M, Neefjes J. 2000. The major substrates for TAP invivo are derived from newly synthesized proteins. Nature 404:774–778. doi:10.1038/35008103

Robinson J, Guethlein LA, Cereb N, Yang SY, Norman PJ, Marsh SGE, Parham P. 2017. Distinguishing functional polymorphism from random variation in the sequences of >10,000 HLA-A, -B and -C alleles. Plos Genet 13:e1006862. doi:10.1371/journal.pgen.1006862

Rock KL, Reits E, Neefjes J. 2016. Present Yourself! By MHC Class I and MHC Class II Molecules. Trends Immunol 37:724–737. doi:10.1016/j.it.2016.08.010

Schuster H, Shao W, Weiss T, Pedrioli PGA, Roth P, Weller M, Campbell DS, Deutsch EW, Moritz RL, Planz O, Rammensee H-G, Aebersold R, Caron E. 2018. A tissue-based draft map of the murine MHC class I immunopeptidome. Sci Data 5:180157. doi:10.1038/sdata.2018.157

Serwold T, Gonzalez F, Kim J, Jacob R, Shastri N. 2002. ERAAP customizes peptides for MHC class I molecules in the endoplasmic reticulum. Nature 419:480–483. doi:10.1038/nature01074

She X, Rohl CA, Castle JC, Kulkarni AV, Johnson JM, Chen R. 2009. Definition, conservation and epigenetics of housekeeping and tissue-enriched genes. BMC Genomics 10:269. doi:10.1186/1471-2164-10-269

Shen XZ, Billet S, Lin C, Okwan-Duodu D, Chen X, Lukacher AE, Bernstein KE. 2011. The carboxypeptidase ACE shapes the MHC class I peptide repertoire. Nat Immunol 12:1078–1085. doi:10.1038/ni.2107

Shen XZ, Lukacher AE, Billet S, Williams IR, Bernstein KE. 2008. Expression of Angiotensin-converting Enzyme Changes Major Histocompatibility Complex Class I Peptide Presentation by Modifying C Termini of Peptide Precursors*. J Biol Chem 283:9957–9965. doi:10.1074/jbc.m709574200

Siepel A, Bejerano G, Pedersen JS, Hinrichs AS, Hou M, Rosenbloom K, Clawson H, Spieth J, Hillier LW, Richards S, Weinstock GM, Wilson RK, Gibbs RA, Kent WJ, Miller W, Haussler D. 2005. Evolutionarily conserved elements in vertebrate, insect, worm, and yeast genomes. Genome Res 15:1034–1050. doi:10.1101/gr.3715005

Siepel A, Haussler D. 2004. Combining Phylogenetic and Hidden Markov Models in Biosequence Analysis. J Comput Biol 11:413–428. doi:10.1089/1066527041410472

Söllner JF, Leparc G, Hildebrandt T, Klein H, Thomas L, Stupka E, Simon E. 2017. An RNA-Seq atlas of gene expression in mouse and rat normal tissues. Sci Data 4:170185. doi:10.1038/sdata.2017.185

Subramanian A, Tamayo P, Mootha VK, Mukherjee S, Ebert BL, Gillette MA, Paulovich A, Pomeroy SL, Golub TR, Lander ES, Mesirov JP. 2005. Gene set enrichment analysis: A knowledge-based approach for interpreting genome-wide expression profiles. Proc National Acad Sci 102:15545–15550. doi:10.1073/pnas.0506580102

Sun K, Zhao Y, Wang H, Sun H. 2014. Sebnif: An Integrated Bioinformatics Pipeline for the Identification of Novel Large Intergenic Noncoding RNAs (lincRNAs) - Application in Human Skeletal Muscle Cells. Plos One 9:e84500. doi:10.1371/journal.pone.0084500

Towne CF, York IA, Neijssen J, Karow ML, Murphy AJ, Valenzuela DM, Yancopoulos GD, Neefjes JJ, Rock KL. 2005. Leucine Aminopeptidase Is Not Essential for Trimming Peptides in the Cytosol or Generating Epitopes for MHC Class I Antigen Presentation. J Immunol 175:6605–6614. doi:10.4049/jimmunol.175.10.6605

Tran NH, Qiao R, Xin L, Chen X, Liu C, Zhang X, Shan B, Ghodsi A, Li M. 2018. Deep learning enables de novo peptide sequencing from data-independent-acquisition mass spectrometry. Nat Methods 16:63–66. doi:10.1038/s41592-018-0260-3

Trentini DB, Pecoraro M, Tiwary S, Cox J, Mann M, Hipp MS, Hartl FU. 2020. Role for ribosome-associated quality control in sampling proteins for MHC class I-mediated antigen presentation. Proc Natl Acad Sci USA 117:4099–4108. doi:10.1073/pnas.1914401117

Tscharke DC, Croft NP, Doherty PC, Gruta NLL. 2015. Sizing up the key determinants of the CD8(+) T cell response. Nat Rev Immunol 15:705–716. doi:10.1038/nri3905

Verteuil D de, Granados DP, Thibault P, Perreault C. 2012. Origin and plasticity of MHC I-associated self peptides. Autoimmun Rev 11:627–635. doi:10.1016/j.autrev.2011.11.003

Verteuil D de, Muratore-Schroeder TL, Granados DP, Fortier M-H, Hardy M-P, Bramoullé A, Caron E, Vincent K, Mader S, Lemieux S, Thibault P, Perreault C. 2010. Deletion of immunoproteasome subunits imprints on the transcriptome and has a broad impact on peptides presented by major histocompatibility complex I molecules. Mol Cell Proteomics 9:2034–2047. doi:10.1074/mcp.m900566-mcp200

Villani A-C, Sarkizova S, Hacohen N. 2018. Systems Immunology: Learning the Rules of the Immune System. Annu Rev Immunol 36:813–842. doi:10.1146/annurev-immunol-042617-053035

Vizcaíno JA, Kubiniok P, Kovalchik K, Ma Q, Duquette JD, Mongrain I, Deutsch EW, Peters B, Sette A, Sirois I, Caron E. 2020. The Human Immunopeptidome Project: a roadmap to predict and treat immune diseases. Mol Cell Proteomics 19:31–49. doi:10.1074/mcp.r119.001743

Wadman M, Couzin-Frankel J, Kaiser J, Matacic C. 2020. A rampage through the body. Sci New York N Y 368:356–360. doi:10.1126/science.368.6489.356

Wang D, Eraslan B, Wieland T, Hallström B, Hopf T, Zolg DP, Zecha J, Asplund A, Li L, Meng C, Frejno M, Schmidt T, Schnatbaum K, Wilhelm M, Ponten F, Uhlen M, Gagneur J, Hahne H, Kuster B. 2019. A deep proteome and transcriptome abundance atlas of 29 healthy human tissues. Mol Syst Biol 15:e8503. doi:10.15252/msb.20188503

Xing Y, Hogquist KA. 2012. T-Cell Tolerance: Central and Peripheral. Cold Spring Harb Perspect Biol 4:a006957. doi:10.1101/cshperspect.a006957

Yewdell JW, Reits E, Neefjes J. 2003. Making sense of mass destruction: quantitating MHC class I antigen presentation. Nat Rev Immunol 3:952–961. doi:10.1038/nri1250

York IA, Mo AXY, Lemerise K, Zeng W, Shen Y, Abraham CR, Saric T, Goldberg AL, Rock KL. 2003. The Cytosolic Endopeptidase, Thimet Oligopeptidase, Destroys Antigenic Peptides and Limits the Extent of MHC Class I Antigen Presentation. Immunity 18:429–440. doi:10.1016/s1074-7613(03)00058-x

Zeng J, Liu S, Zhao Y, Tan X, Aljohi HA, Liu W, Hu S. 2016. Identification and analysis of house-keeping and tissue-specific genes based on RNA-seq data sets across 15 mouse tissues. Gene 576:560–570. doi:10.1016/j.gene.2015.11.003

Zhu J, He F, Hu S, Yu J. 2008. On the nature of human housekeeping genes. Trends Genet 24:481–484. doi:10.1016/j.tig.2008.08.004

